# Enhancing non-local interaction modeling for ab initio biomolecular calculations and simulations with ViSNet-PIMA

**DOI:** 10.64898/2026.03.18.712561

**Authors:** Taoyong Cui, Zihan Wang, Tong Wang

**Affiliations:** State Key Laboratory of Membrane Biology Beijing Frontier Research Center for Biological Structure Tsinghua-Peking Center for Life Sciences, School of Life Sciences, Tsinghua University, 100084, Beijing, China

## Abstract

AI-based molecular dynamics simulation brings ab initio calculations to biomolecules in an efficient way, in which the machine learning force field (MLFF) locates at the central position by accurately predicting the molecular energies and forces. Most existing MLFFs assume localized interatomic interactions, limiting their ability to accurately model non-local interactions, which are crucial in biomolecular dynamics. In this study, we introduce ViSNet-PIMA, which efficiently learns non-local interactions by physics-informed multipole aggregator (PIMA) and accurately encodes molecular geometric information. ViSNet-PIMA outperforms all state-of-the-art MLFFs for energy and force predictions of different kinds of biomolecules and various conformations on MD22 and AIMD-Chig datasets, while adapting the PIMA blocks into other MLFFs further achieves 55.1% performance gains, demonstrating the superiority of ViSNet-PIMA and the universality of the model design. Furthermore, we propose AI^**2**^BMD-PIMA to incorporate ViSNet-PIMA into AI^**2**^BMD simulation program by introducing “Transfer Learning-Pretraining-Finetuning” scheme and replacing molecular mechanics-based non-local calculations among protein fragments with ViSNet-PIMA, which reduces AI^**2**^BMD’s energy and force calculation errors by more than 50% for different protein conformations and protein folding and unfolding processes. ViSNet-PIMA advances ab initio calculation for the entire biomolecules, amplifying the application values of AI-based molecular dynamics simulations and property calculations in biochemical research.

## Main

Biomolecular dynamics simulation is a fundamentally computational approach for life sciences research[1, 2]. The simulations make sense only if they are efficient and accurate[3–6]. Classical molecular dynamics simulation, grounded in molecular mechanics (MM), adopts pre-defined functions to calculate the molecular potential energy. Despite their speed, MM methods usually fall short in describing certain quantum effects, limiting their calculation accuracy[7–9]. As a comparison, quantum molecular dynamics simulation provides ab initio calculations by using quantum mechanics (QM)[10], while these methods such as density functional theory (DFT) have difficulty scaling to large biomolecular systems due to the prohibitively computational costs[11].

Most recently, by employing the machine learning force field (MLFF), AI^2^BMD[12], for the first time, transforms protein dynamics simulations to ab initio accuracy with the computational cost orders of magnitude lower than DFT. It adopts a generalizable protein fragmentation scheme to divide various proteins into commonly used protein fragments and employs the equivariant graph neural network (EGNN) ViSNet[13] as the MLFF. ViSNet is trained on molecular data calculated at DFT level, predicts the potential energy and atomic forces for protein units and then drives the molecular movements during the simulations. Although AI^2^BMD achieves ab initio calculation accuracy for protein fragments, the non-local interactions among protein fragments are still calculated by MM, i.e., Coulomb force and van der Waals, limiting ab initio calculations on protein backbones instead of the entire biomolecules. Whereas non-local interactions play key roles in protein folding, conformational changes and bioactivities[14–16], accurately modeling non-local interactions by MLFF is vital for characterizing biomolecular dynamics.

MLFF has emerged as a rapidly advancing field, thanks to incorporating equivariant features regarding to the SO(3) rotation group in the graph neural networks. ViSNet[13], Equiformer V2[17], Mace[18] and other state-of-the-art EGNNs[19–24] have significantly improved MLFF’s prediction performance and data usage efficiency[21]. However, most existing MLFFs are designed based on the assumption that interatomic interactions are localized[25]. This limitation arises from the intrinsic model design, which aggregates information only from the atoms within a predefined radius threshold in EGNN. Although this design is highly effective at capturing local interactions, it is far insufficient to accurately model non-local interactions, particularly those arising from electrostatics[26, 27]. Furthermore, the inaccessible cost to generate adequate DFT data for different conformations of the entire large biomolecules poses a significant challenge for training MLFFs, making it difficult to learn the non-local interactions[28].

To tackle the challenge, Ewald message passing[29] leverages a Fourier space framework with frequency-based cutoffs, rooted in the Ewald summation approach, to capture non-local interactions, while ViSNet-LSRM [30] leverages the breaking of retrosynthetically interesting chemical substructures (termed as “BRICS”) fragmentation algorithm [31] and fragmentation-based message passing to explicitly incorporate non-local interactions. However, these methods often ignore the anisotropic electron density distributions arising from different conformations of the same molecule, limiting their generalizability, particularly for larger biomolecules and rare conformations. This limitation undermines their applicability in AI-based molecular dynamics simulations, where accurately capturing phenomena such as electrostatic polarization is critical for describing biomolecular behavior. Therefore, incorporating directional information to better represent anisotropic electron density remains an open challenge in the development of MLFFs.

In this study, we transfer the physical theory into model design for accurately modeling non-local interactions. Multipole expansion is a widely used physical approach to depict molecular electron density distributions as the combinations and interactions among point charges, dipoles, quadrupoles, and higher-order terms[32]. Given the coordinates of a molecular conformation, this approach calculates and updates the global multipole information iteratively and then can accurately characterize the non-local electrostatic effects till calculation convergence [32, 33]. Inspired by multipole expansion theory, we designed Physics-Informed Multipole Aggregator (PIMA), which learns molecular non-local interactions by iteratively updating each node’s equivariant features in EGNNs. Then, we incorporated the PIMA module into ViSNet as ViSNet-PIMA, which learns and encodes both local and non-local interactions simultaneously. We made comprehensive evaluations on the widely used biomolecular DFT datasets, MD22[34] consisting five different kinds of biomolecules two kinds of supramolecules and AIMD-Chig[35] consisting of two million different conformations of the protein Chignolin. ViSNet-PIMA surpassed all state-of-the-art MLFFs on energy and force predictions. Furthermore, when adapting PIMA into different MLFFs, all deep learning models with PIMA outperformed the same models but with other non-local interaction modeling algorithms for all test cases, showing that PIMA could serve as a universal and effective component in EGNNs for non-local interaction modeling. Notably, when explicitly evaluated the non-covalent interactions for different dimer systems, ViSNet-PIMA exhibited much more consistent predictions with DFT than the original version of ViSNet with empirical Coulomb and van der Waals functions. Furthermore, ViSNet-PIMA can accurately characterize the non-local electrostatic forces in anion-cation systems and biomolecular association process in an aqueous environment.

More importantly, to enhance ab initio calculation for the entire protein during AI-based molecular dynamics simulations, we proposed AI^2^BMD-PIMA that adapted ViSNet-PIMA into AI^2^BMD to replace MM calculations for non-local and interfragment interactions among protein units. To achieve it, we innovatively proposed a “Transfer Learning—Pretraining—Finetuning” scheme in AI^2^BMD to train ViSNet-PIMA by transferring the hidden representations embedded in protein units into model inputs, pretraining the model with massive data on MM level and finetuning it with only a minimal amount of DFT data. With the highly efficient data usage strategy, AI^2^BMD-PIMA thoroughly achieved ab initio calculations for the entire proteins by remarkably reducing the energy and force calculation errors by more than 50% for different proteins and showing much more accurate calculations during protein folding and unfolding simulations and thermodynamics property estimations when compared to the original version of AI^2^BMD, paving the way for accurately studying protein dynamics mediated by non-local interactions and amplifying its application values in biochemical fields.

### Overview of ViSNet-PIMA and AI^2^BMD-PIMA

The architecture of ViSNet-PIMA (Fig. 1 a) integrates geometric deep learning with physics-informed message passing mechanisms, enabling accurate predictions of energies and forces while preserving 3D equivariance. This is achieved by incorporating the PIMA module into EGNNs (e.g., ViSNet) and jointly training them to model both local and non-local interactions simultaneously. The short-long interactions are first learnt and represented by ViSNet layers. Then by leveraging multipole expansion and iterative feature updates, PIMA aggregates non-local information from two phases, i.e., “Field Response” phase and “Interaction Integration” phase. In the “Field Response” phase, PIMA utilizes dipole-dipole responses to model electrostatic polarization effects. Dipole embeddings are initialized by the atomic equivariant features from ViSNet layers and iteratively updated and refined through polarization responses to external electric fields and environmental modulation from the neighboring atoms. This process mimics the physical induction of molecular dipoles toward convergence, while implicitly encoding many-body interactions. Furthermore, the iterative updates and refinements enable progressive convergence to stable dipole configurations for each atom, which is critical for non-local effect modeling. In the “Interaction Integration” phase, the invariant atomic features are enhanced through dipole interaction modeling. The updated dipole embeddings drive feature transformations that combine non-local electrostatic effects with local chemical environments. These augmented features, now encoding both local and non-local interactions, are subsequently aggregated to predict atomic contributions to the molecular potential energy.

**Figure 1:**
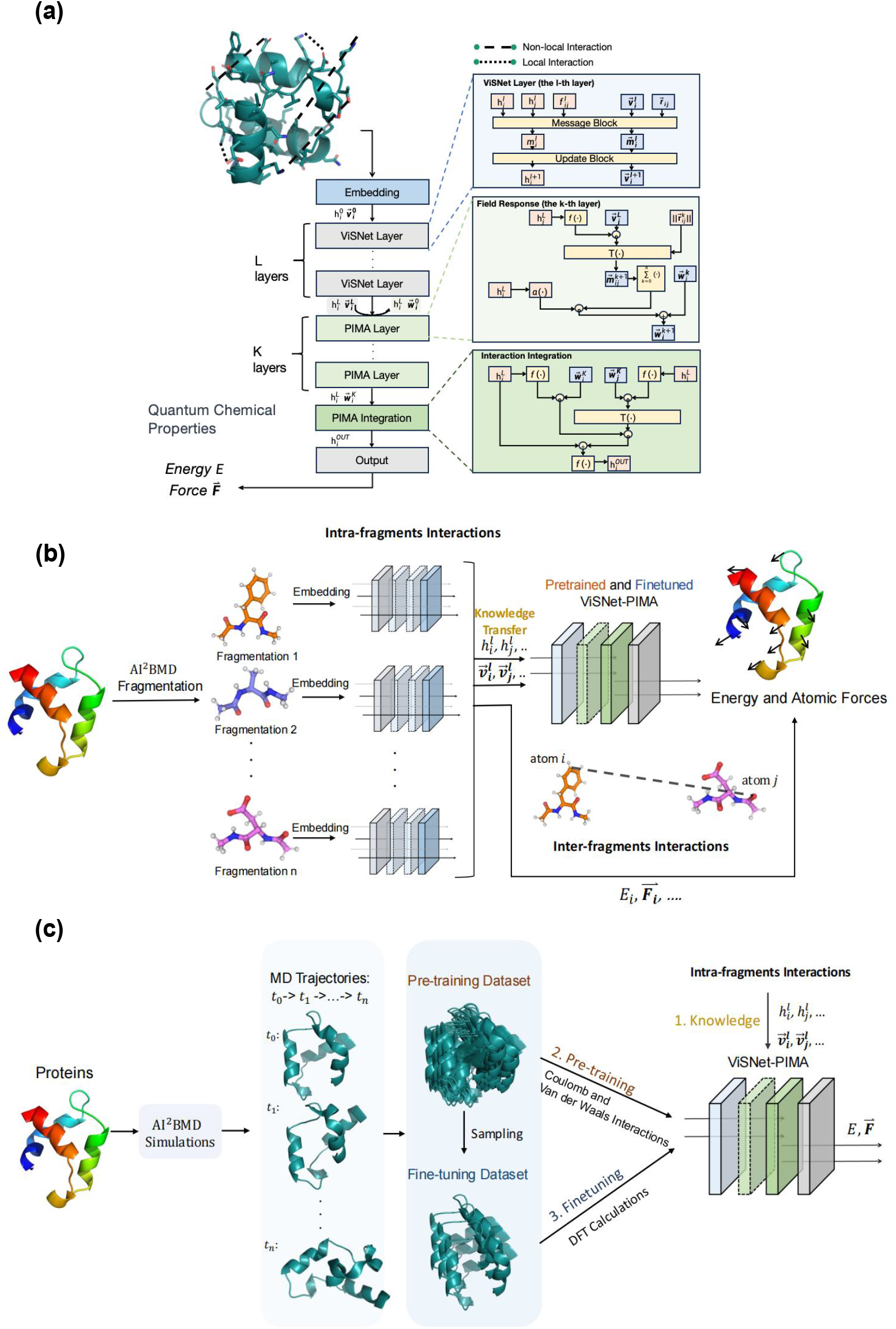
The overall pipeline of ViSNet-PIMA and its incorporation into AI^2^BMD. **a**. The model architecture of ViSNet-PIMA. It takes the three-dimensional structure of the molecule as input, extracts geometric information via a series of ViSNet layers and PIMA layers and predicts quantum chemical properties, such as energies and forces by the output layer. In ViSNet-PIMA, ViSNet layers are used to learn and update scalar features 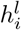 and vectorized features 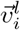 through equivariant message passing. The inputs of the first PIMA layer are the relative position 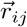 and the outputs of the *L*-th ViSNet layer, including the vectorized features 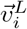, represented as 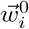 and the scalar features 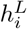. In the “Field Response” phase, the vectorized feature 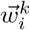 is iteratively updated through a gated residual connection of a series of PIMA layers. In the “Interaction Integration” phase, 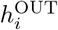 is calculated by applying the Hadamard product to the aggregated vectorized messages, performing Cartesian interaction calculations, and incorporating a gated residual connection. Besides ViSNet-PIMA, the PIMA module can be adapted into other EGNNs to form EGNN-PIMA for accurate molecular modeling. **b**. The architecture of AI^2^BMD-PIMA. Proteins are divided into protein units by AI^2^BMD fragmentation process. The potential energy and atomic forces of each protein unit are calculated. Then through a “Transfer Learning-Pretraining-Finetuning” scheme, ViSNet-PIMA transfers knowledge on intra-fragments interactions within protein units as model inputs and learns inter-fragments interactions among protein units by pretraining and finetuning processes. **c**. The “Transfer Learning-Pretraining-Finetuning” scheme. Through AI^2^BMD simulations, samples in MD trajectories are evenly extracted to construct the pretraining dataset, followed by a random sampling to create a much smaller finetuning dataset. MM calculations, i.e., Coulomb and van der Waals are performed on the pretraining dataset for non-local interactions among protein units, while DFT calculations are performed on the finetuning dataset to get the residual energies and forces between the entire molecules and those summed from protein units. ViSNet-PIMA leverages the transferred knowledge on local interactions as part of the inputs, learns non-local interactions from the massive samples in the pretraining dataset, and finetunes the model parameters on the finetuning dataset calculated at DFT level. Ultimately, this scheme enables ViSNet-PIMA to predict energy and forces related to inter-fragments interactions with ab initio accuracy.

In this study, we further present the incorporation of ViSNet-PIMA into AI^2^BMD, i.e., AI^2^BMD-PIMA to achieve ab initio calculations for the entire molecules and its applications during simulations. The simulation system leverages a fragmentation process to divide proteins into smaller fragments, enabling to model local (intra-fragment) and non-local (inter-fragment) interactions, respectively. As illustrated in Fig. 1 b, the framework employs a hierarchical modeling strategy: local intra-fragment interactions are firstly captured by ViSNet, while non-local inter-fragment interactions are then modeled via ViSNet-PIMA. A key design lies in the “Transfer LearningPretraining-Finetuning” scheme (Fig. 1 c), where pretraining on Coulomb and van der Waals interactions of the conformations sampled from AI^2^BMD simulation trajectories efficiently encodes fundamental physical principles. This process enables subsequent finetuning with minimal DFT data to achieve high-fidelity predictions. The scheme not only enhances computational efficiency but also maintains the high precision required for studying complicated biomolecular systems, offering a scalable solution for ab initio molecular simulations.

### Energy and force predictions by ViSNet-PIMA

We first evaluated ViSNet-PIMA’s performance on energy and force predictions by taking both different kinds of biomolecules and different conformations of the same protein into consideration. The MD22 dataset comprises MD trajectories for seven distinct molecular systems, including proteins, lipids, carbohydrates, nucleic acids, and supramolecules. These systems, ranging from 42 to 370 atoms, are designed to challenge MLFFs with their complexity and diversity. To ensure consistency, the data split for MD22 dataset follows the methodology outlined in [30]. This dataset allows us to rigorously evaluate ViSNet-PIMA’s performance across various molecular types and sizes. In parallel, the AIMD-Chig dataset provides a focused assessment of various conformations of the protein. It comprehensively samples the conformational space of Chignolin, a 166-atom mini-protein, serving as a benchmark for handling the dynamic and diverse conformations of full-atom proteins.

We compared ViSNet-PIMA with state-of-the-art algorithms, including PaiNN[36], Equiformer v2[17], So3krates[37], Mace[18], and ViSNet[13] on MD22 dataset, as shown in Table 1. ViSNet-PIMA achieved the smallest MAEs on both energy and force predictions for all 7 molecules, demonstrating the prominent advantages of ViSNet-PIMA over ViSNet and other state-of-the-art models. Although So3krates is specifically designed to describe non-local quantum mechanical effects across arbitrary length scales, ViSNet-PIMA outperformed it on all molecules. Since Equiformer v2 is not an energy-conservation model, it exhibits large discrepancies in energy and force predictions especially for the more complex supramolecules, implying its potential limitations in accurately capturing energy dynamics.

**Table 1:** Mean absolute errors (MAE) of energy (kcal/mol) and force (kcal/(mol·Å)) predictions for 5 kinds of biomolecules and 2 kinds of supramolecules on MD22 dataset. The best one in each category is highlighted in bold.

| Molecule |  | PaiNN | SO3krates | Equiformer v2 | Mace | ViSNet | ViSNet-PIMA |
| --- | --- | --- | --- | --- | --- | --- | --- |
| Ac-Ala3-NHMe | energy | 0.0955 | 0.3372 | 0.4344 | 0.0620 | 0.0901 | **0.0530** |
|  | forces | 0.1598 | 0.2443 | 0.0821 | 0.0876 | 0.0799 | **0.0744** |
| DHA | energy | 0.1811 | 0.3792 | 0.1192 | 0.1317 | 0.0938 | **0.0784** |
|  | forces | 0.1666 | 0.2424 | 0.0732 | 0.0646 | 0.0618 | **0.0534** |
| Stachyose | energy | 0.3200 | 0.4425 | 0.2523 | 0.1244 | 0.1351 | **0.0843** |
|  | forces | 0.2842 | 0.4357 | 0.1128 | 0.0821 | 0.0934 | **0.0677** |
| AT-AT | energy | 0.2012 | 0.1785 | 0.3081 | 0.1093 | 0.1309 | **0.0772** |
|  | forces | 0.2464 | 0.2169 | 0.2067 | 0.0992 | 0.0960 | **0.0781** |
| AT-AT-CG-CG | energy | 0.3767 | 0.3456 | 0.7572 | 0.1578 | 0.2032 | **0.1571** |
|  | forces | 0.3321 | 0.3325 | 0.2545 | 0.1153 | 0.1293 | **0.1142** |
| Buckyball catcher | energy | 0.4523 | 0.4327 | 1.4532 | 0.2720 | 0.5020 | **0.2032** |
|  | forces | 0.4047 | 0.3823 | 0.1669 | 0.1844 | 0.1844 | **0.0935** |
| Double-walled nanotube | energy | 0.6134 | 0.5577 | 3.1772 | 0.6326 | 0.8002 | **0.4522** |
|  | forces | 0.3573 | 0.3986 | 0.2452 | 0.3421 | 0.3621 | **0.2243** |

Through the entire MD22 dataset, ViSNet-PIMA consistently outperformed the vanilla ViSNet, achieving an average improvement of 39 % in energy predictions and 19% in force predictions. Furthermore, for the supramolecules buckyball catcher and double-walled nanotube, ViSNet-PIMA achieved remarkable improvements in accuracy against ViSNet, with energy predictions improved by 44%-55% and force predictions improved by 38%-49%. These results highlight ViSNet-PIMA’s ability to effectively model non-local interactions, which is essential for accurately describing complex supramolecular systems. In addition, when compared to the second-best performing models, ViSNet-PIMA maintains a substantial lead, with average improvements of 15.2% in energy predictions and 12.8% in force predictions.

For AIMD-Chig dataset, as shown in Figure 2 a-b and Supplementary Table S1, ViSNet-PIMA achieved a 13% error reduction in energy prediction compared to the second-best model PaiNN and a 9% improvement in force prediction accuracy over the suboptimal approach Equiformer v2, suggesting its enhanced capability in distinguishing the tiny chemical property differences among different conformations. To further investigate ViSNet-PIMA’s performance on folded, unfolded and intermediate states of the protein, we plotted Chignolin’s free energy landscape based on time-lagged independent component analysis[38] and picked up some representative conformations for analysis (Figure 2 c). As shown on the free energy landscape, ViSNet-PIMA outperformed ViSNet for the unfolded state as the conformation is more relaxed. For the folded states, both models exhibited comparable performance. Interestingly, for the transition states between the folded and unfolded conformations, ViSNet-PIMA outperformed ViSNet on energy prediction by a large margin. We infer that this observation may be resulted from the non-local electrostatic interactions in protein folding and unfolding processes, which are especially prominent in the transition states.

**Figure 2:**
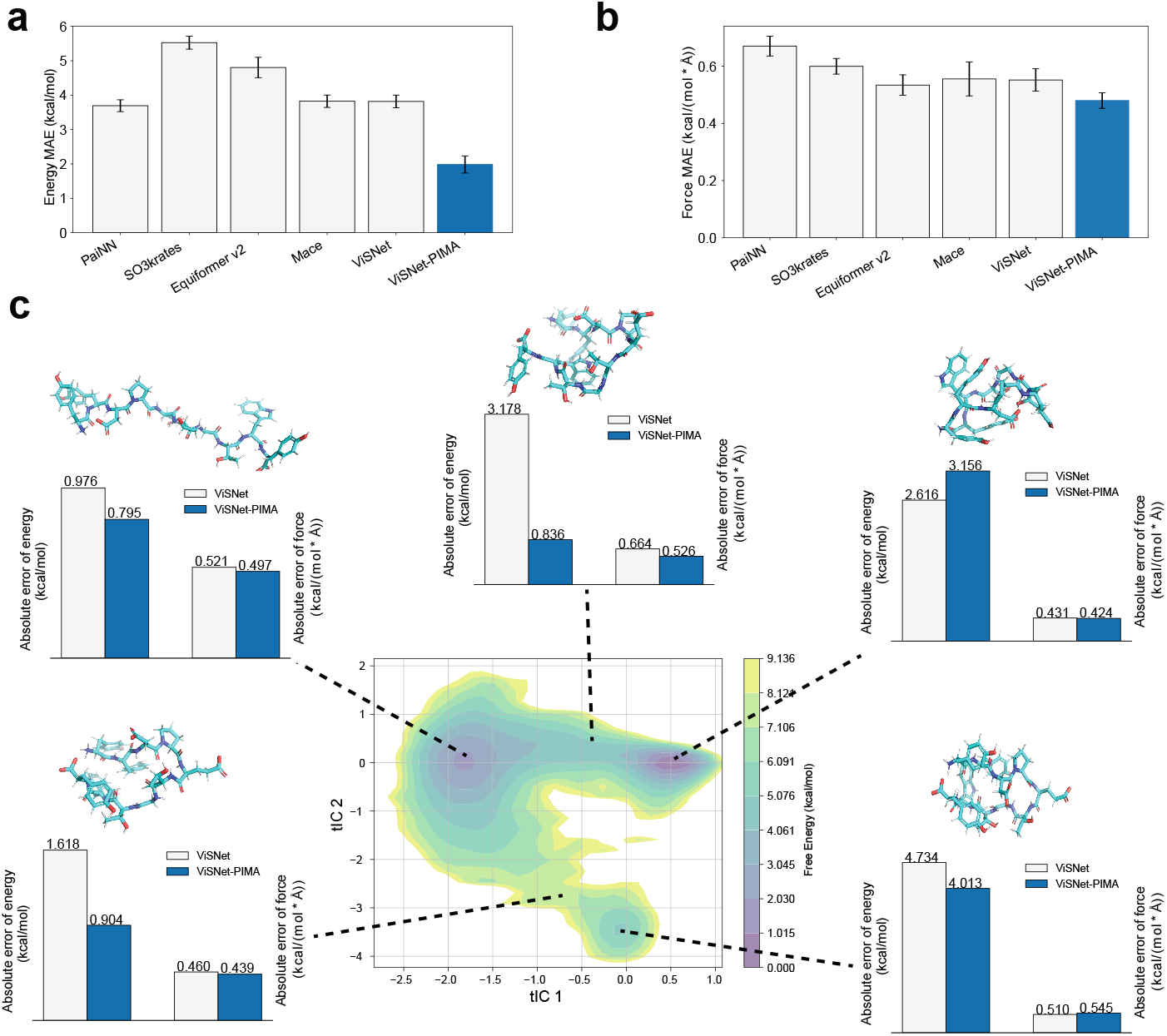
Evaluations of ViSNet-PIMA on AIMD-Chig dataset. **a**. Mean absolute error of energy on AIMD-Chig predicted by ViSNet-PIMA and other MLFFs. **b**. Mean absolute error of forces on AIMD-Chig predicted by ViSNet-PIMA and other MLFFs. In **a** and **b**, the prediction made by ViSNet is shown in the blue bar, while those made by other MLFFs are shown in light grey bars. The standard error for each prediction is also shown. **c**. Comparison of ViSNet and ViSNet-PIMA on representative conformations of Chignolin on the free energy landscape. Five representative structures with two folded, one unfolded and two intermediates states are drawn with sticks. For each structure, the energy and force predictions by ViSNet and ViSNet-PIMA are shown with light grey bars and blue bars, respectively. The values of absolute energy and force errors are given on the top of the corresponding bars.

In addition, we also examined ViSNet-PIMA’s performance on dipole prediction. This evaluation was made by training on the molecular dipole moments on the dataset in [39]. When evaluated on the test set, ViSNet-PIMA achieved a Mean Absolute Error (MAE) of 0.000368 a.u., a much higher accuracy than 4.4-fold improvement over the baseline ViSNet (with a MAE of 0.001627 a.u.). This remarkable improvement demonstrates that PIMA’s physics-informed model design can learn the physical representation of the molecular charge distribution well and thus facilitates molecular property prediction, such as energy, force and dipole. All the evaluation results demonstrate ViSNet-PIMA’s versatility, excelling in diverse molecular systems, while also accurately modeling the intricate dynamics and conformational changes during the protein dynamics simulations.

### Universality and effectiveness of the PIMA module

To further examine the universality and effectiveness of the PIMA module, we chose two EGNN model architectures, i.e., PaiNN and ViSNet as the base models and adapted different non-local interaction modeling algorithms, including PIMA, Ewald and LSRM into them and benchmarked their performance on the MD22 dataset. As shown in Table 2, both PaiNN-PIMA and ViSNet-PIMA demonstrate conspicuous performance enhancements, with improvements reaching up to 55% over the corresponding base models, highlighting PIMA’s universal applicability across diverse EGNN frameworks. We further compared PIMA with Ewald Message Passing[29] and LSRM[30] and found that both PaiNN and ViSNet models with PIMA surpassed those with either Ewald or LSRM for energy and force predictions on all MD22 molecules. For example, a reduction of 61% energy MAE and 23% force MAE in stachyose is observed made by ViSNet-PIMA compared to ViSNet-Ewald, while PaiNN-PIMA also demonstrates universal performance enhancements, slashing Double-Walled Nanotube force MAE by 63% compared to PaiNN-LSRM. These results further highlight that PIMA can act as a universal module to be adapted into MLFFs and excels in modeling non-local interatomic interactions.

**Table 2:** Mean absolute errors (MAE) of energy (kcal/mol) and force kcal/(mol·Å)) predicted by MLFFs with Ewald, LSRM and PIMA modules on MD22 dataset. The performace of the base models PaiNN and ViSNet are shown in Table 1. The best one in each category is highlighted in bold.

| Molecule |  | PaiNN-Ewald | PaiNN-LSRM | PaiNN-PIMA | ViSNet-Ewald | ViSNet-LSRM | ViSNet-PIMA |
| --- | --- | --- | --- | --- | --- | --- | --- |
| Ac-Ala3-NHMe | energy | 0.1272 | 0.0931 | 0.0823 | 0.1475 | 0.0748 | **0.0530** |
|  | forces | 0.1671 | 0.1662 | 0.1411 | 0.0840 | 0.0761 | **0.0744** |
| DHA | energy | 0.1778 | 0.1332 | 0.1203 | 0.0824 | 0.1195 | **0.0784** |
|  | forces | 0.1664 | 0.1297 | 0.1221 | 0.0708 | 0.0717 | **0.0534** |
| Stachyose | energy | 0.2522 | 0.1756 | 0.1642 | 0.2144 | 0.1055 | **0.0843** |
|  | forces | 0.2821 | 0.2398 | 0.1811 | 0.0875 | 0.0767 | **0.0677** |
| AT-AT | energy | 0.1923 | 0.1042 | 0.0827 | 0.1688 | 0.1007 | **0.0772** |
|  | forces | 0.2232 | 0.1887 | 0.1139 | 0.1070 | 0.0881 | **0.0781** |
| AT-AT-CG-CG | energy | 0.3361 | 0.2431 | 0.1998 | 0.2428 | 0.2447 | **0.1571** |
|  | forces | 0.2924 | 0.3221 | 0.2012 | 0.1479 | 0.1293 | **0.1142** |
| Buckyball catcher | energy | 0.3523 | 0.6599 | 0.3176 | 0.2799 | 0.4220 | **0.2032** |
|  | forces | 0.4023 | 0.4912 | 0.3723 | 0.1178 | 0.1069 | **0.0935** |
| Double-walled nanotube | energy | 0.5927 | 1.9442 | 0.5542 | 0.7944 | 1.8223 | **0.4522** |
|  | forces | 0.4095 | 0.9199 | 0.3375 | 0.2833 | 0.3683 | **0.2243** |

We then investigated how unique design in PIMA contributes to model’s performance gains by conducting ablation experiments on double-walled nanotube, since the supramolecule is the largest and most challenging one in MD22 dataset. The results, summarized in Supplementary Table S2, reveal critical insights into the elaborate model design and performance gains of PIMA. First, ViSNet-PIMA with six ViSNet layers followed by three PIMA layers outperformed the original version of ViSNet model with nine ViSNet layers, despite identical total number of model layers. The result demonstrates that explicitly modeling non-local interactions via physics-guided layers is more effective than simply increasing the model capacity for local interactions. Notably, directly expanding the receptive field of the ViSNet layer by increasing the neighboring atom threshold to 100Å, i.e., ViSNet-L layers, caused out-of-memory failures, underscoring PIMA’s computational efficiency in decoupling interaction range from memory costs. Secondly, reversing the order of ViSNet layers and PIMA layers in the model decreased the prediction accuracy, indicating that the local atomic environment representation learning should precede molecular non-local interaction modeling. This mirrors the physical hierarchy where local quantum effects dictate local polarization, which subsequently influences non-local electrostatic fields. Thirdly, the variant PIMA-S by updating both the invariant and equivariant features simultaneously exhibited much poorer performance than PIMA that only updates equivariant features in the “Field Response” phase. Keeping invariant features fixed while updating equivariant features in PIMA’s “Field Response” phase is similar with the physical calculation process in polarizable force field, that the point charges are fixed while the induced dipoles are iteratively updated.

Notably, despite the extra non-local representation capacity, a PIMA layer is distinctly more lightweight than a ViSNet layer, containing approximately 0.5 million parameters compared to ViSNet’s 1.1 million parameters. We benchmarked the model training speed on the Ac-Ala3-NHMe molecule from the MD22 dataset. As shown in Supplementary Table S3, a 6-layer ViSNet model trains at 110 s/epoch. ViSNet-PIMA model, which has 9 layers (6 ViSNet + 3 PIMA) in total, is trained at 143 s/epoch. For a fair comparison against a model with the same depth, we also trained a 9-layer large ViSNet model, which required 159 s/epoch. In addition, a state-of-the-art model, MACE is also compared. It consumed 1,024 s/epoch during training, about 7.16 times slower than ViSNet-PIMA. These results indicate that while the 9-layer ViSNet-PIMA is slower than a smaller 6-layer model, it is notably faster than ViSNet of the same depth. This is due to the parameter efficiency of the PIMA layers.

In addition, we also designed ViSNet and ViSNet-PIMA with fewer layers and estimated the model size, training cost and performance compared with other state-of-the-art models. The small version of ViSNet consists of 3 ViSNet layers with 128 features in each layer, while the small version of ViSNet-PIMA consists of 2 ViSNet layers and 3 PIMA layers with 128 features in each layer. As shown in Supplementary Table S4, even with much fewer parameters, the small version of ViSNet-PIMA still achieved the highest prediction accuracy among all machine learning force fields when evaluated on the molecule AT-AT-CG-CG in MD22 dataset. Although MACE has fewer model parameters than ViSNet-PIMA (small), its model training cost is higher than 10 times of that of ViSNet-PIMA and its accuracy is lower than ViSNet-PIMA. Such results further consolidate that the accuracy gains from PIMA are not merely a product of increased computational load, but stem from a more efficient, physically-informed architecture and thus can provide one of the best performance-to-cost tradeoffs.

### Non-local interaction learning and representation

To evaluate ViSNet-PIMA’s ability to model non-local interactions, we adopted the dimer molecules from SPICE dataset[40, 41], in which the structures originate from the benchmark data in DES370K[42]. These dimer molecules consist of separate monomers at varying distances, and the interaction energies between the seperate monomers are primarily dominated by non-covalent interactions. Thus, the dimer dataset serves as a suitable benchmark for probing MLFFs’ capabilities in non-local interaction learning[43, 44].

The dimer dataset consists of 3,490 pairs of dimers, with 80 % selected for training and validation, and the remaining 20 % reserved as the test set to avoid data leakage. We then trained ViSNet-PIMA, which includes 2 ViSNet layers and 3 PIMA layers with a threshold set to 3 Å, on the training set. For comparison, we also evaluated a 5-layer ViSNet under identical settings. In these systems, ViSNet-PIMA achieved an energy error of 1.29 kcal/mol, while ViSNet exhibited a higher energy error of 2.81 kcal/mol. The integration of PIMA with ViSNet resulted in over 50 % error reduction in non-local interaction modeling, highlighting the importance of explicitly accounting for non-local interactions in model design.

Furthermore, we examined ViSNet-PIMA’s qualitative behavior in modeling noncovalent interactions compared to original ViSNet by selecting representative dimer systems in detail. As shown in Figure 3, the chosen dimers represent interactions involving cations, anions, charged compounds, and polar compounds, effectively serving as test cases for charge-dipole and dipole-dipole interactions. Additional examples are provided in Supplementary Figure S1. Compared to ViSNet, ViSNet-PIMA successfully recovered the binding energy curves of separate monomers in all representative systems. While once the distance between two monomers exceeds the threshold of the ViSNet layer, ViSNet model failed to capture energy fluctuations, even with additional message passing layers. We also analyzed the relative energy errors for the representative systems shown in Figure 3. For all four test dimer systems, ViSNet-PIMA maintains a low relative error in the longer-range distances (Supplementary Figure S2). In contrast, the baseline ViSNet model exhibits an increasing relative error as the separation distance increases beyond the local interactions. This limitation arises from how molecular graphs are constructed in local message-passing EGNNs. Specifically, these models build molecular graphs based on a predefined atom neighboring threshold, treating separated monomers as distinct graphs that cannot exchange interaction messages when the distance of two monomers exceeds the threshold. Notably, even the distance of two monomers within the atom neighboring threshold, ViSNet failed to characterize the energy curves in some cases. This observation indicates that the proposed PIMA module enhances not only non-local interaction modeling but also facilitates local interaction learning, thereby improving the overall expressive power and generalization ability of MLFFs.

**Figure 3:**
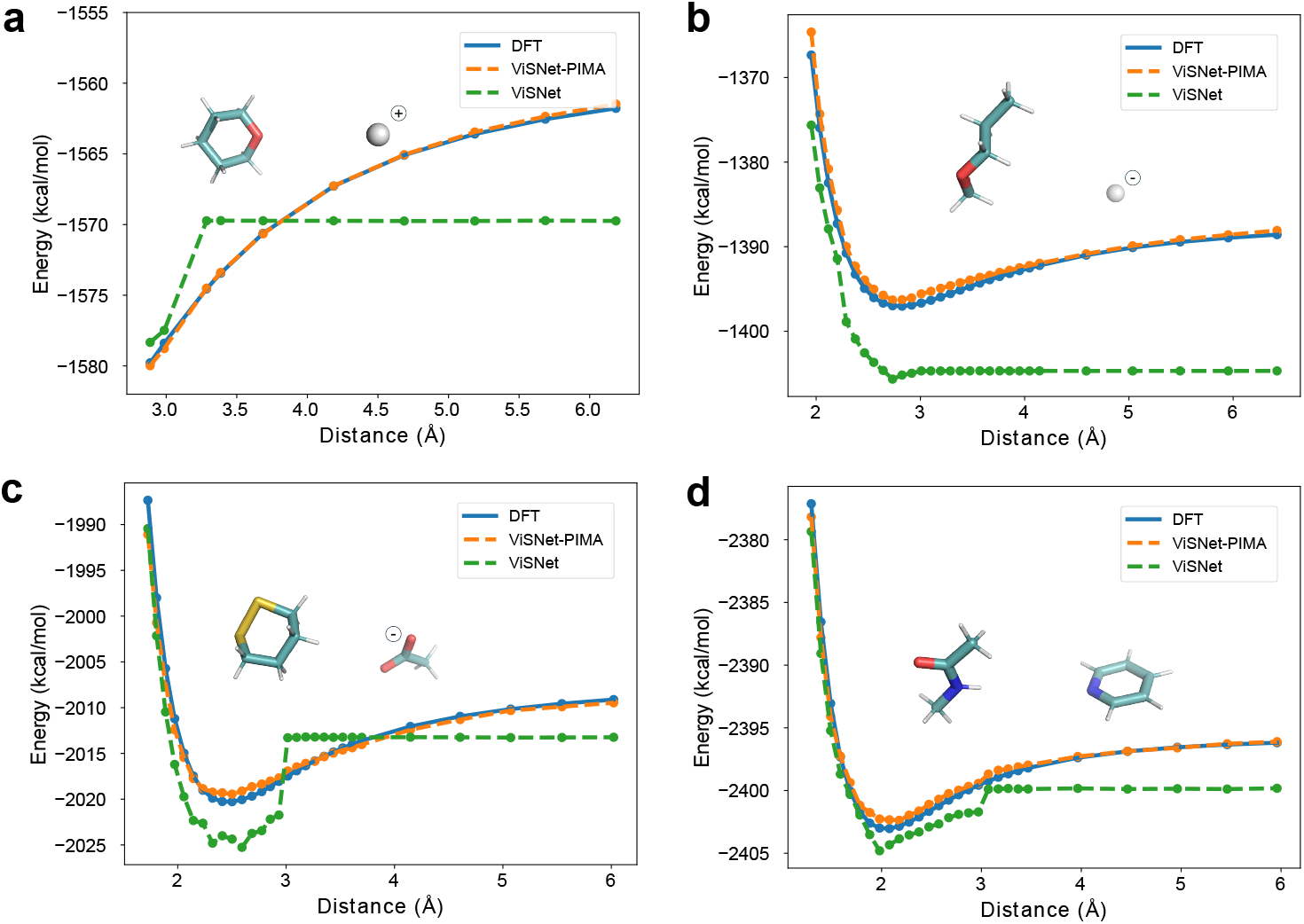
Evaluations of non-local interactions on dimer molecules by ViSNet-PIMA. **a**. The binding energy curves of a dimer between a polar compound and a cation (sodium ion) calculated by DFT, ViSNet-PIMA and ViSNet. **b**. The binding energy curves of a dimer between a polar compound and an anion (chloride ion) calculated by DFT, ViSNet-PIMA and ViSNet. **c**. The binding energy curves of a dimer between a polar compound and a charged compound calculated by DFT, ViSNet-PIMA and ViSNet. **d**. The binding energy curves of a dimer between two polar compounds calculated by DFT, ViSNet-PIMA and ViSNet. In **a**-**d**, the structures of the dimers are also shown.

In addition, we further benchmarked another baseline model by augmenting the local message-passing ViSNet model with an explicit, empirical pairwise correction (i.e., Coulomb and van der Waals) for non-local interactions. We evaluated the performance of the new baseline (termed as “ViSNet-MM”) on the four test systems included in Figure 3, and found this baseline performed distinctly worse than ViSNet-PIMA and failed to correctly describe the potential energy surfaces. As shown in Supplementary Figure S3 a and b, the MM corrections of Coulomb and van der Waals failed to model the non-local decay of the separate dimers. Furthermore, as shown in Supplementary Figure S3 c and d, the MM corrections also inaccurately captured interactions at very short distances (*r <* 2 Å).. This result exhibits that simple pairwise functions with point charges may be inadequate for these challenging systems, further underscoring the necessity of the learned PIMA module to capture the many-body polarization effects as well as leverage the predictive power of base models.

### MD simulations for various molecules

To evaluate the applicability of ViSNet-PIMA in MD simulations, we first examined ViSNet-PIMA’s energy conservation by performing a MD simulation for Ac-Ala_3_-NHMe molecule from the MD22 dataset. The system was simulated for 10,000 simulation steps with 1 fs per step under NVE ensemble driven by the ViSNet-PIMA potential. Over the entire 10 ps trajectory, compared to the total energy of −6.2 ∗10^5^ kcal/mol, the fluctuation of the total energy was observed to be less than 1 kcal/mol (*<* 0.002% of the total energy, Supplementary Figure S4 a). Furthermore, as shown in Supplementary Figure S4 b-d, the forces on each axis are also counterbalanced. This negligible energy drift confirms that our model generates forces consistent with the energy, leading to stable dynamics.

We then leveraged ViSNet-PIMA as MLFF to perform 100 ps MD simulations for three different kinds of molecules on MD22 dataset, including Ac-Ala_3_-NHMe, stachyose, and the buckyball catcher. The simulations were performed by ASE simulation framework[45] at 300K under NVT ensemble in vacuum. We analyzed the relative potential energy fluctuations along the simulation trajectories and compared them with DFT calculations. As shown in Supplementary Figure S5 a–c, ViSNet-PIMA exhibited consistent energy calculation values with DFT for all three molecules, demonstrating its ab initio accuracy in capturing potential energy fluctuations throughout the simulation trajectories.

We evaluated the interatomic distance distributions of the three kinds of molecules derived from the simulation trajectories. As shown in Supplementary Figure S5 d–f, the protein, carbohydrate and supramolecue dominant interactions at different length scales, resulting in distinct distance distribution profiles. From Ac-Ala_3_-NHMe to the buckyball catcher, the interatomic distance distributions exhibit a shift towards longer distances as the molecular size increases. For the buckyball catcher, the interactions between the buckyball and the catcher are purely non-covalent, dominated by *π*–*π* interactions, which are inherently non-local ones. Therefore, accurately describing such interactions is crucial for the reasonable behavior during MD simulations[46]. During the 100 ps simulation driven by ViSNet-PIMA, the buckyball remained inside the catcher, indicating that our model can effectively capture such non-local interactions and thus describe the dynamic motions of the supramolecule with high fidelity. In addition, we also analyzed the kinetic properties from the simulation trajectories by examining the vibrational spectra of these molecules (Supplementary Figure S5 g–i). The spectra were calculated as the Fourier transform of the velocity autocorrelation function derived from the trajectories. Our results were able to reproduce the characteristic vibrational peaks associated with each kind of molecule, such as the C=C stretching (∼ 1600 *cm*^−1^) and C-H stretching (∼ 3000 *cm*^−1^) of buckyball catcher[34]. The experimental findings further highlight the exceptional performance of ViSNet-PIMA in closely aligning with DFT calculations, underscoring its robust capability in modeling both local and non-local interactions. This precision in capturing molecular dynamics across varying scales not only validates the model’s accuracy but also demonstrates its ability to integrate local model expressiveness with non-local effects.

We then designed a system where electrostatic polarization effects are particularly prominent to further examine the performance gains of ViSNet-PIMA. The system consists of a guanidinium cation and an acetate anion simulated in vacuum to cleanly isolate the non-local electrostatic effects. This serves as a simple test system for one of the most important interactions in biology: the salt bridge between the side chains of Arginine (Arg) and Aspartate (Asp). The formation of such salt bridges is a key driving force in protein folding, structural stability, and molecular recognition. We performed three independent 1 ns molecular dynamics simulations for each approach: AI driven simulations by ViSNet-PIMA and one of the most recent state-of-the-art AI driven simulation program MACE-OFF[47] for comparison. The simulations started with the ion pair at an initial separation of 12 Å.

The results in Supplementary Figure S6 show a distinct and qualitative difference in the dynamical behavior sampled by the two approaches, providing clear evidence for the necessity of the PIMA module. As shown in all three independent simulation trajectories sampled by MACE-OFF, the two ions did not show any trend to close to each other and even drifted further apart over the course of the 1 ns simulation, indicating that by employing the local message-passing model, MACE-OFF failed to capture the non-local electrostatic attraction. In contrast, ViSNet-PIMA successfully and robustly captured the non-local attraction. In all three replicas, the ions rapidly associated within the first about 150 ps to form a stable contact ion pair. The ion pair kept stably bound in the remaining simulation time, fluctuating around the expected equilibrium distance. Such results further consolidate ViSNet-PIMA can effectively capture non-local and electrostatic polarization-dominant interactions.

Furthermore, ViSNet-PIMA is designed to be capable of handling solvated systems as it allows water molecules as inputs. To quantitatively validate its performance in the presence of water, following the procedures in MACE-OFF[47], we have benchmarked ViSNet-PIMA trained on the SPICE dataset containing different kinds of molecules as well as water. An independent water molecule subset in SPICE dataset consisting of 1-50 water molecules was used as the test set. Note that there is no sample overlapped between the training and test sets to avoid data leakage. We compared ViSNet-PIMA with MACE-OFF, and the results shown in Supplementary Figure S7 indicate that ViSNet-PIMA achieved a distinctly lower prediction error on the water test set compared to MACE-OFF. This demonstrates the robustness and accuracy of ViSNet-PIMA in describing systems where explicit solute-water interactions are present. To further examine ViSNet-PIMA’s performance in handling solvent in MD simulations, we have implemented a jax-version ViSNet-PIMA for efficient MD simulations on larger systems. The model was trained on the SPICE dataset and deployed it into ASE simulation framework. We then performed simulations in an explicit solvent with umbrella sampling and calculated the Potential of Mean Force (PMF) for the association of two p-Cresol molecules, where p-Cresol is a common minimalist model for the Tyrosine side chain. As shown in Supplementary Figure S8, the resulting PMF shows a clear free energy minimum corresponding to the stable T-shaped *π*-stacking structure of the dimer system. This outcome is in agreement with established chemical principles for the association of aromatic molecules in water, which is driven by a complex interplay between the hydrophobic effect and direct solute-solute interactions. These results indicated that ViSNet-PIMA can not only model non-local interactions but also accurately characterize the biomolecular association process in a realistic aqueous environment.

### The AI^2^BMD-PIMA simulation program

To advance ab initio calculations and full-atom simulations for different proteins, we integrated ViSNet-PIMA into AI^2^BMD[12] via a “Transfer Learning-Pretraining-Finetuning” scheme. AI^2^BMD-PIMA synergistically combines the predictive power of the baseline AI^2^BMD for local protein fragments with ViSNet-PIMA’s proficiency in capturing non-local interactions. This integration reduces the data requirements for non-local interaction modeling and enables efficient learning and representation of molecular interactions across different scales. In AI^2^BMD, ViSNet was initially trained on an extensive dataset of protein units, i.e., Protein Unit Dataset (PUD)[12] to effectively capture local chemical knowledge across diverse environments. The training process enables the model to generate continuous latent atom embeddings, which describe local chemical environments derived from the molecular graph, allowing for a detailed representation of molecular interactions at the atomic level. In “Transfer Learning-Pretraining-Finetuning” scheme, the knowledge acquired by the ViSNet potential in the baseline AI^2^BMD was directly transferred to ViSNet-PIMA. Subsequently, ViSNet-PIMA underwent pretraining on conformations derived from AI^2^BMD simulation trajectories to assimilate empirical non-bonded calculations by Coulomb and van der Waals. Following the pretraining, a limited set of energy and force labels calculated at the DFT level was generated for the target proteins. The pretrained ViSNet-PIMA was then finetuned on the tiny dataset with an energy MAE of 3.73 kcal/mol and a force MAE of 0.58 kcal/(mol·Å). Notably, over 80,000 pretraining samples were derived from the simulated trajectories of four proteins, including Chignolin, Trp-cage, WW domian and albumin-binding domain (ABD), while for each protein, only 1,000 finetuning samples were generated at DFT level. As illustrated in Supplementary Figure S9 and Supplementary Figure S10, the pretraining and finetuning datasets display similar global energy distributions, while also maintaining subtle variations per energy bin. This result consolidates the necessity of our pretrain-finetune framework, i.e., the pretraining process provides general knowledge for molecular representation learning and enhances the model’s generalization ability for different proteins, while the finetuning process further refines and improves the model’s ability to distinguish tiny energy differences among protein conformations. In addition, a comprehensive summary to demonstrate the differences between AI^2^BMD and AI^2^BMD-PIMA is shown in Supplementary Table S5.

To comprehensively evaluate the performance gains of ViSNet-PIMA for AI^2^BMD, we compared the MAE of potential energy and atomic forces calculated by AI^2^BMD-PIMA, the baseline AI^2^BMD and MM respectively, with the DFT calculations as the reference values shown in Figure 4. For Chignolin, Trp-cage, WW domian and ABD, same to the conformations employed in [12], 50 folded, 50 unfolded and 100 intermediate structures were chosen for each protein based on the radius of gyration[48] for evaluation. Compared with MM, AI^2^BMD has already significantly reduced the calculation errors on energy and atomic forces. Its sequence-based fragmentation approach treats all local interactions within protein units of the protein at ab initio level, enabling accurate calculations on backbone atoms. However, the non-local interactions among side-chain atoms and non-adjacent protein fragments are still calculated at MM level, which becomes the main factor that hinders calculation accuracy. By integrating AI^2^BMD-PIMA, such interactions become learnable by our proposed learning scheme. As a result, AI^2^BMD-PIMA exhibited an energy MAE of 0.0309 kcal/mol per atom and force MAE of 0.7138 kcal/(mol·Å) averaged on the four proteins, while the AI^2^BMD exhibited an energy MAE of 0.0677 kcal/mol per atom and force MAE of 1.7807 kcal/(mol·Å). We normalized the energy differences by the number of atoms to directly compare the results across different proteins. For all four proteins, AI^2^BMD-PIMA reduced the calculation errors by over 50% on both energy and forces, showcasing the importance of accurately modeling inter-fragment interactions. We further investigated the detailed performance of AI^2^BMD and AI^2^BMD-PIMA on unfolded, intermediate and folded states of these proteins, respectively. The results are summarized in Figure 4 d-e and Supplementary Figure S11. As shown in Figure 4 d, AI^2^BMD exhibited a large force prediction error for the folded conformations than unfolded and intermediate conformations. This result is observed on all four test proteins. We infer the inconsistent performance among different states of proteins may come from the treatment of non-local terms in AI^2^BMD. To validate it, we defined the non-local interactions in AI^2^BMD as the residual atomic forces between the forces calculated on the whole protein and those calculated from protein units and analyzed the distributions of residual forces among different topologies of the protein. As shown in Supplementary Figure S12, a positive shift of the residual force distributions is captured from the unfolded to the folded state, indicating that such non-local interactions are more distinct in folded states. Since the empirical Coulomb and Van der Waals functions may be inadequate to accurately model them, the baseline AI^2^BMD’s fall short in describing the folded conformations. As a comparison, replacing the MM calculations by ViSNet-PIMA, the updated AI^2^BMD exhibited consistent and ab initio calculation accuracy across different states of proteins, while achieving ordersof-magnitude speed acceleration compared to DFT, as evidenced by Supplementary Figure S13.

**Figure 4:**
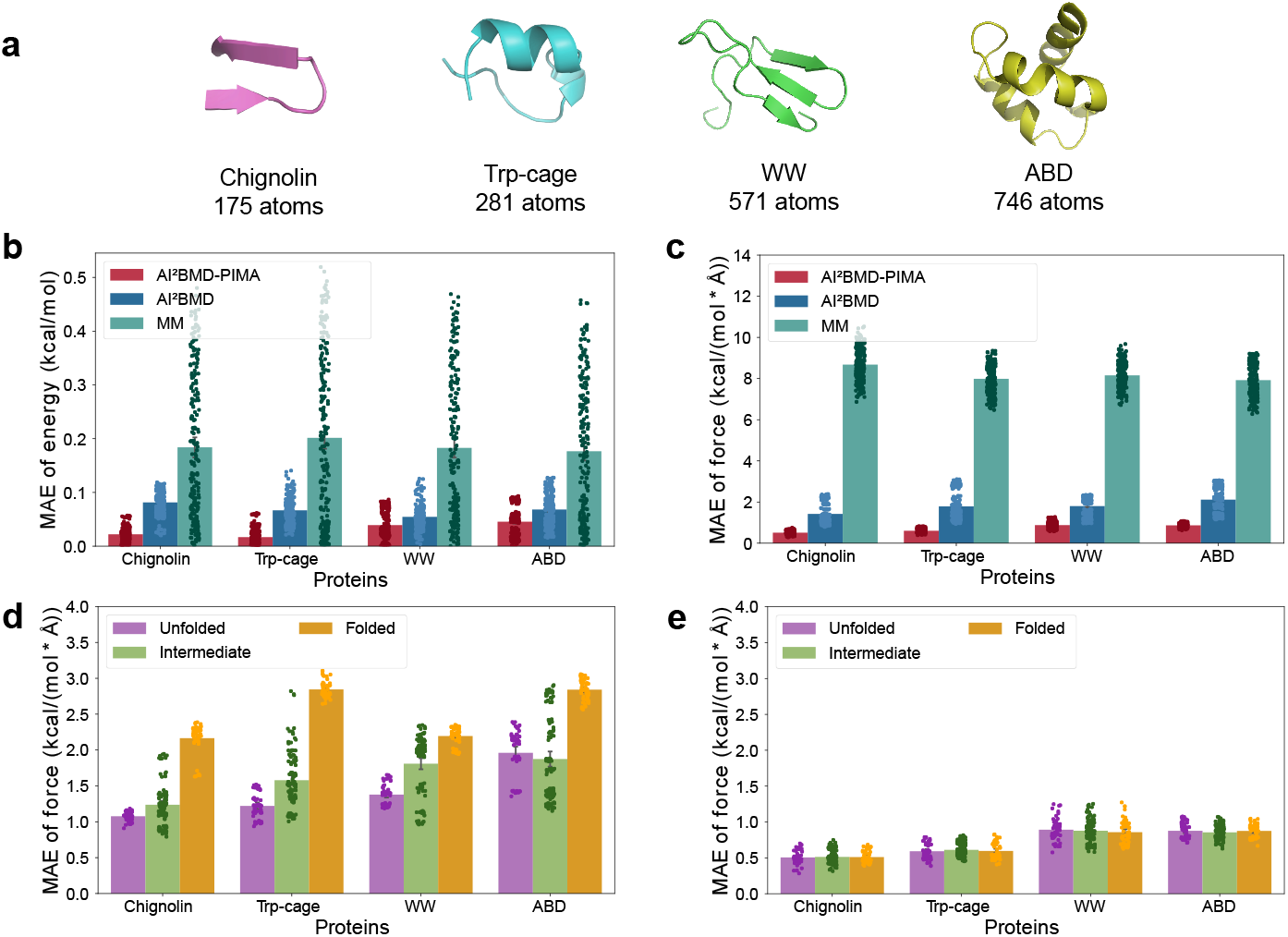
Evaluations of AI^2^BMD-PIMA on energy and force calculations for different proteins. **a**. Folded structures of the four evaluated proteins. The structures are shown in cartoon. **b**. Mean absolute error (MAE) of energy per atom calculated by AI^2^BMD-PIMA, AI^2^BMD and MM respectively. For MM calculations, the potential energy of each structure has that of a reference structure subtracted. **c**. Mean absolute error of force calculated by AI^2^BMD-PIMA, AI^2^BMD and MM. **d**. Mean absolute error of force calculated by AI^2^BMD on unfolded, intermediate and folded structures, respectively. **e**. Mean absolute error of force calculated by AI^2^BMD-PIMA on unfolded, intermediate and folded structures, respectively. In **b**-**e**, the DFT calculation act as the reference values. The absolute energy or force error compared to DFT for each sample is also shown in scatter.

Furthermore, we analyzed the fluctuations in energy and force during the protein folding and unfolding processes. Starting from a folded and an unfolded conformation of Chignolin, 10ns simulations were performed by AI^2^BMD with a 10Å water box at 300K under NVT ensemble. After simulations, a total of 200 conformations for each trajectory were selected at even intervals, and the protein structures were extracted for energy and force evaluations. As illustrated in Fig 5, for the protein folding process, the MAE of energy and force predicted by AI^2^BMD-PIMA were 0.0308 kcal/mol per atom and 1.0667 kcal/(mol·Å), respectively. As a comparison, those predicted by AI^2^BMD without ViSNet-PIMA were 0.1570 kcal/mol per atom and 2.3065 kcal/(mol·Å). Similar results can be seen on the protein unfolding process as the MAE of energy and force made by AI^2^BMD-PIMA were 0.0171 kcal/mol per atom and 0.8204 kcal/(mol·Å), compared with those made by AI^2^BMD without ViSNet-PIMA are 0.1469 kcal/mol per atom and 1.9566 kcal/(mol·Å). Notably, AI^2^BMD exhibits smaller MAE values during the second half unfolding process and the first half of folding process compared to the corresponding counterparts, indicating the discrepancy of force predictions for different conformations illustrated in Fig 4 d. In contrast, AI^2^BMD-PIMA shows consistent prediction accuracy during the folding and unfolding processes, aligning with the results in Fig 4 e, further demonstrating its values and usefulness in modeling protein folding in a more accurate manner. In addition, we also evaluated the simulation speed of AI^2^BMD-PIMA. As shown in Supplementary Table S6, the baseline AI^2^BMD, which uses classical potentials for inter-fragment interactions, runs at approximately 8 seconds per 100 simulation steps for simulating the protein Trp-cage. The new AI^2^BMD-PIMA runs at 10 seconds per 100 steps. In contrast, employing the one of the most recent state-of-the-art AI driven simulation program MACE-OFF[47] to perform MD simulation consumes 27 seconds per 100 steps, much slower than AI^2^BMD-PIMA. This result indicates a reasonable trade-off to capture non-local interactions at ab initio accuracy with moderate computational overhead.

**Figure 5:**
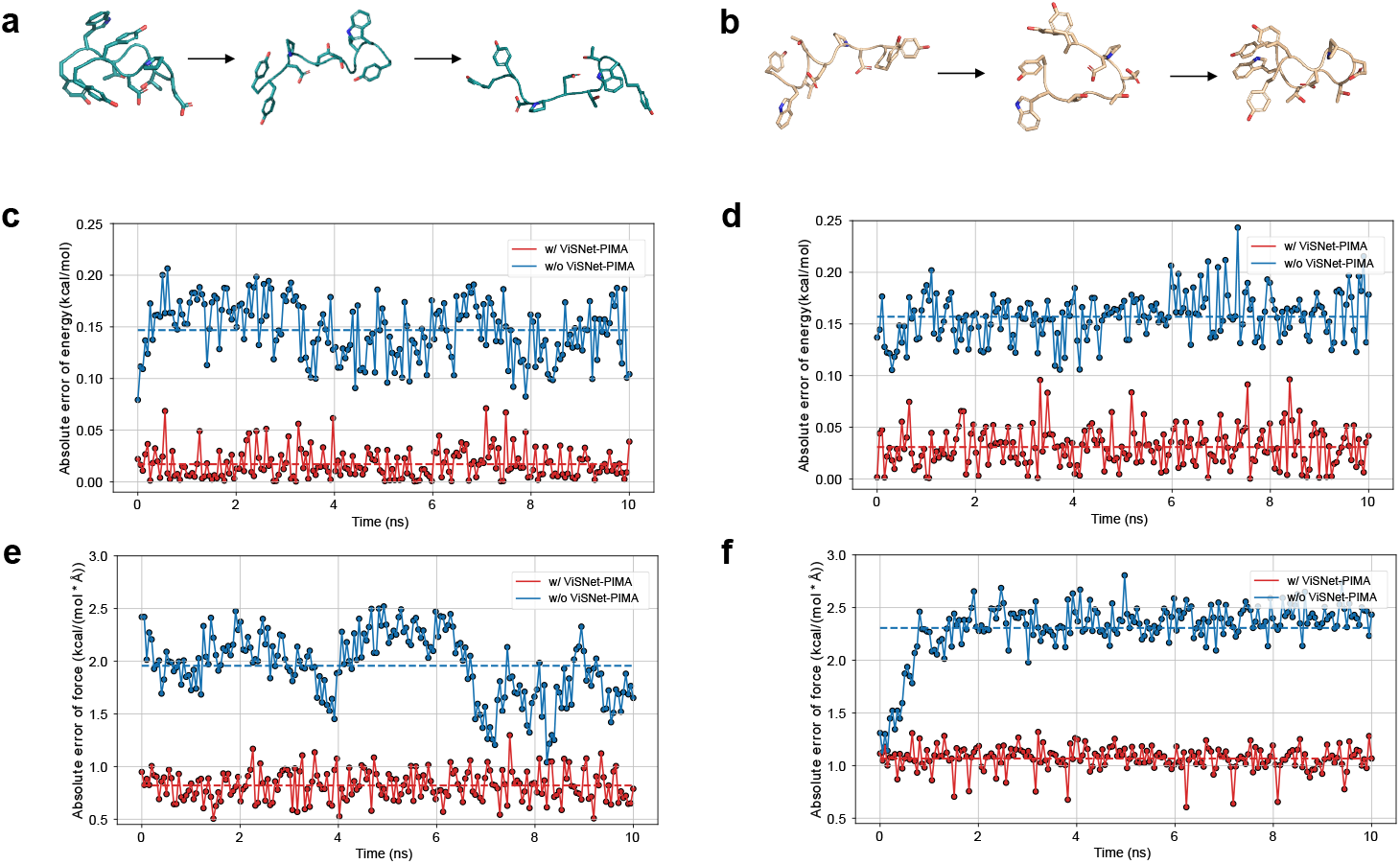
Evaluations of energy and force calculations for Chignolin folding and unfolding simulation processes. **a, b**. Representative structures along the 10ns simulation trajectories performed by AI^2^BMD starting from a folded state (a) and an unfolded state (b). **c, d**. Absolute errors of potential energy per atom of the protein calculated by AI^2^BMD-PIMA and those calculated by AI^2^BMD during the simulations for the unfolding process shown in **a** and the folding process shown in **b. e, f**. Absolute errors of force of the protein calculated by AI^2^BMD-PIMA and those calculated by AI^2^BMD during the simulations for the unfolding process shown in **a** and the folding process shown in **b**. In **c**-**f**, the red and blue dash lines indicate the MAE of energy and force calculations by AI^2^BMD-PIMA and AI^2^BMD during the whole simulations, respectively.

Beyond energy and force evaluations during simulations, we further examined the usefulness of AI^2^BMD-PIMA in thermodynamics property estimations. Following thesame procedures adopted in AI^2^BMD [12], we estimated folding free energy differences (Δ*G*) and melting temperatures (*T*_*m*_) for three representative proteins (WW, BBA, and Protein G) by classifying folded/unfolded ensembles using the Q-score and reweighting potential energy distributions via AI^2^BMD-PIMA. As shown in Figure 6, compared with AI^2^BMD, AI^2^BMD-PIMA achieved consistent improvements in *T*_*m*_ prediction accuracy, with error reductions of 29% for WW, 13% for BBA, and 10% for Protein G. For Δ*G* estimation, AI^2^BMD-PIMA delivered improvements of 17% for BBA and 14% for Protein G, while maintaining a slightly higher accuracy for WW. These results indicate that AI^2^BMD-PIMA not only retains the *ab initio*level accuracy of AI^2^BMD, but also systematically enhances thermodynamic property prediction, particularly for *β*-rich and mixed *α/β* proteins.

**Figure 6:**
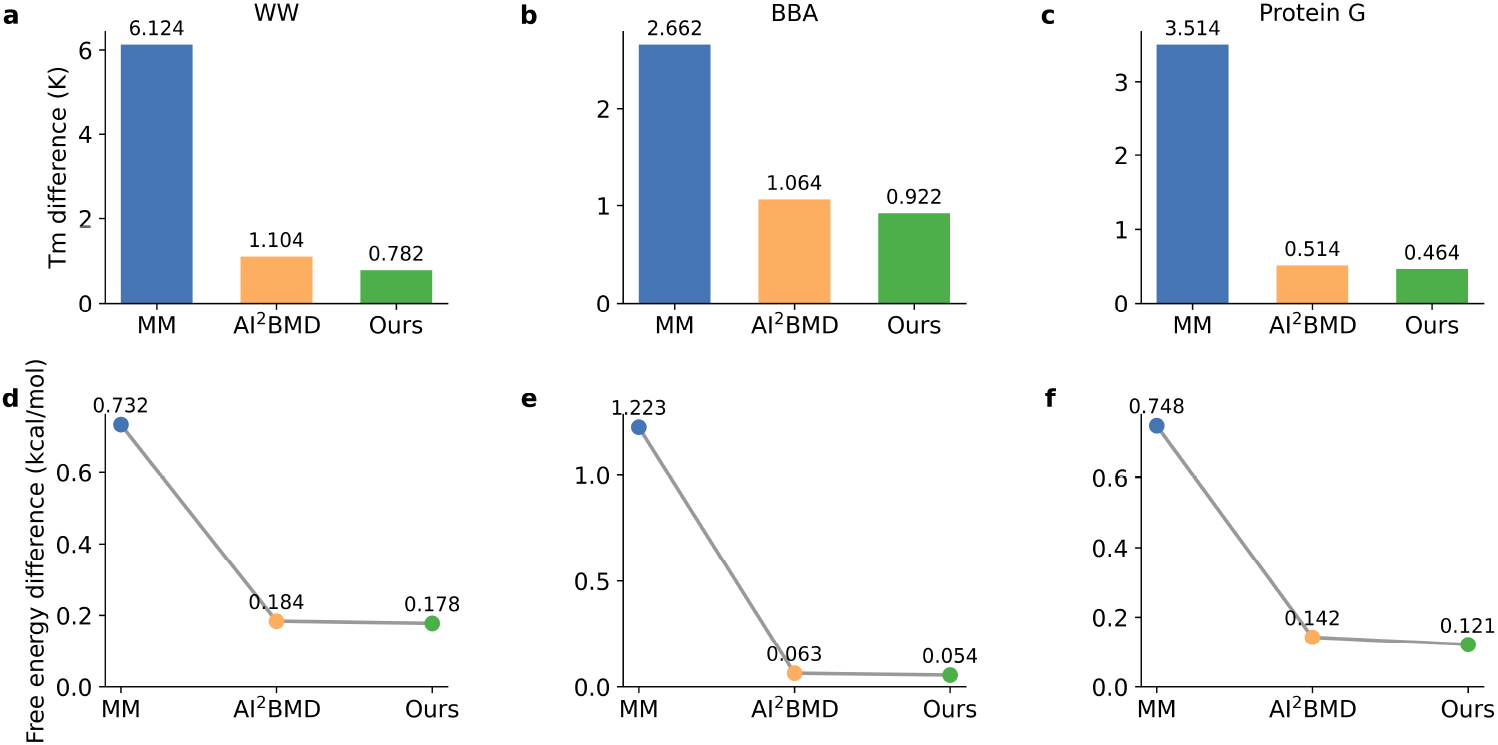
Evaluations of AI^2^BMD-PIMA (termed as”Ours”), AI^2^BMD and MM on melting temperature and folding free energy estimations for three proteins, WW, BBA and protein G. (**a-c**.) The absolute differences of melting temperature (*T*_*m*_) for WW (**a**), BBA (**b**), and Protein G (**c**) estimated by MM, AI^2^BMD, and AI^2^BMD-PIMA. (**d-f**.) The absolute differences of folding free-energy (Δ*G*) for WW (**d**), BBA (**e**), and Protein G (**f**) estimated by MM, AI^2^BMD, and AI^2^BMD-PIMA. The performance of AI^2^BMD and MM are derived from AI^2^BMD paper.

## Discussions

In this study, we present ViSNet-PIMA, which exemplifies the integration of physical priors with machine learning force fields to enhance non-local interaction modeling and outperforms state-of-the-art models by a large margin, when benchmark evaluated on different kinds of biomolecules and conformations.

Non-local interaction modeling is a long-standing challenge in MLFF design, where recent progress can be broadly grouped into three paradigms: (i) local models augmented with explicit classical terms (e.g., Coulomb or van der Waals corrections), (ii) local models with learned non-local corrections via extended message passing or multipolar responses (e.g., SpookyNet[26], Latent Ewald Summation[49], LSRM[30]), and (iii) global-descriptor-based models such as sGDML[34] and AIMNet[50]. ViSNet-PIMA falls into the second category, introducing a multipole-inspired iterative dipole refinement that mimics polarization convergence, thereby improving both physical fidelity and generalization.

The prominent performance gains of ViSNet-PIMA dominantly attribute to the Physics-Informed Multipole Aggregator (PIMA). By effectively combining multipole expansion into model design, PIMA iteratively updates each node’s equivariant features, enabling efficient aggregation of non-local information and capturing electrostatic polarization effects. As PIMA is adopted into different models and exhibits consistently accurate energy and force predictions, we could prospect PIMA will act as a universal and non-local interaction modeling approach, be incorporated into various deep neural networks for molecular geometric modeling and play key roles in chemical, biological and material fields.

By introducing ViSNet-PIMA into AI^2^BMD, it achieves ab initio calculations for the entire proteins. AI^2^BMD has addressed the localized quantum effects through fragment-based approach, while its reliance on MM descriptions for interactions among fragments is inadequate to capture non-local electronic polarization and non-local quantum interactions. Our AI^2^BMD-PIMA has overcome the barriers by integrating geometric deep learning with physics-informed machine learning force fields, which expands the ab initio calculation from protein backbone to the entire protein, specifically for side chains and non-consecutive protein fragments and thus will provide more valuable perspectives for studying protein dynamics mediated by non-local interactions in the future research.

One of the notable design in AI^2^BMD-PIMA is the “Transfer Learning—Pretraining—Finetuning” scheme. By transferring the hidden representations learnt from protein units, ViSNet-PIMA naturally acquires the local chemical environment information of the atoms and focuses on the non-local interaction learning among protein units. The transfer learning strategy lowers the difficulty of molecular representation learning and improves model training and inference efficiency with a relatively small model of ViSNet-PIMA. Furthermore, instead of directly training models on the DFT dataset, the combination of pretraining and finetuning processes with both MM and DFT calculations greatly reduces the DFT data generation consumption, alleviating the dilemma of the prohibitive cost to generate adequate DFT data for different conformations of the entire protein. In addition, although AI^2^BMD-PIMA has achieved ab initio calculation for the entire proteins, acquiring DFT data for larger biological systems remains a challenge in computational chemistry. To address this limitation, the future studies will explore advanced strategies to minimize DFT computational demands while maintaining accuracy, such as enhanced sampling to efficiently explore conformational space, active learning to strategically select structures, unsupervised learning with unlabeled data and parameter fitting from wet-lab experiments. Despite the simulations of two p-Cresol molecules in solvent system as a proof-of-concept example, it is still challenging to perform simulations on larger solvated systems due to scability issues. Following AI^2^BMD, AI^2^BMD-PIMA still treats the solvent using a classical polarizable force field. However, our framework is designed to be flexible enough to adopt emerging fragmentation schemes that can extend quantum mechanical treatment to solute-solvent interactions. Such subsequent investigations will be applied for larger biomolecular systems, particularly those involving complex allosteric interactions or multi-domain protein dynamics, which bridges the gap between quantum mechanical accuracy and biomolecular simulation scalability, with potential applications in enzyme engineering, rational drug design and biomechanism detection.

## Methods

### Preliminaries

#### Graph neural networks

In machine learning force fields (MLFFs), the input molecules are represented as graphs *G* = (*V, E*), where *V* denotes atoms, and *E* denotes edges, mainly based on interatomic distances *r*_*ij*_. Atomic features *h*_*i*_ are initialized based are their atomic numbers and then iteratively refined by graph neural networks through the message passing mechanism. In the *l* th message passing layer, the message that atom *i* receives from its neighboring atom *j* can be defined as follows, where *M* denotes a learnable function that governs the message computation and *α*_*ij*_ denotes the interactions between *i* and *j*:

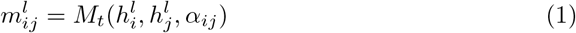

For atom *i*, its features are then updated by a learnable update function *U* to aggregate all the messages from *i*’s neighbors:

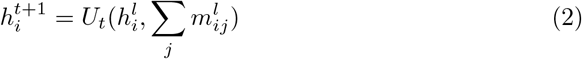

#### Equivariance

A function *f* : *X* → *Y* is said to be equivariant with respect to a group *G*, if for any element *g* of *G*:

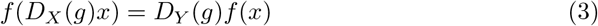

where *D*_*X*_ and *D*_*Y*_ denote the group representations in the vector space *X* and *Y*, respectively. In particular, if *D*_*Y*_ = *I* for all g, *f* is said to be invariant. When predicting potential energy and atomic forces based on atomic positions, MLFFs are required to be equivariant with regard to the operations of *SO*(3) group[51]. This type of equivariance acts as an inductive bias for machine learning models, leading to the development of equivariant GNNs (EGNNs). In EGNNs, each atom possesses invariant features *h*_*i*_, vectorized features 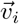, or higher order equivariant features. Each message passing layer utilizes both invariant and equivariant features together. The equivariant features are updated as follows:

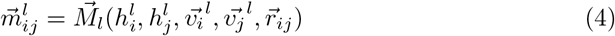

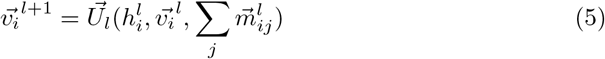

where 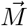 and 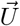 are required to preserve equivariance.

#### Multipole expansion

Considering that an electrostatic charge distribiution is *ρ*(*r*), its electric potential 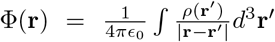 can be approximated by a sum of multipole moments, derived from the Taylor series expansion of 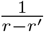 [33]. By defining the point charge 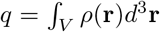 as the monopole moment, 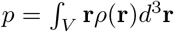 as the dipole moment, the electric potential can be written in the following multipole expansion formula:

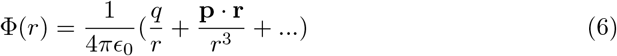

Some polarizable force fields utilize the theory of multipole expansion to model electrostatic interactions between atoms in molecular systems[52]. In these force fields, each atom is assigned with a point charge and a dipole moment. The dipole moment can be induced by external electrostatic fields and will be iteratively updated till convergence. The electrostatic interactions between atoms can be interpreted as the interactions between point charges and dipole moments based on multipole expansion.

#### Overview of the original version of AI^*2*^BMD

AI^2^BMD employs a universal protein fragmentation approach, dividing proteins into 21 kinds of protein units[12]. These units are then reassembled to compute the overall potential energy and atomic forces. A DFT dataset for all kinds of protein units, i.e., the Protein Unit Dataset (PUD) was built by comprehensively exploring the conformational space of such fragments and calculating at DFT level[12]. The PUD dataset was employed to train the equivariant GNN, ViSNet[13] as the AI^2^BMD potential. AI^2^BMD adopted a hybrid calculation strategy for the simulation system. The energies (*E*_prot_) and forces 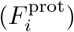 of the entire protein are calculated as linear combinations of those from the corresponding fragments (*E*^prot_units^ and 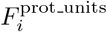), while interactions among fragments are assessed using the classical Coulomb equation and Lennard-Jones potential, as follows:

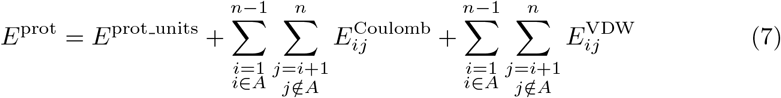

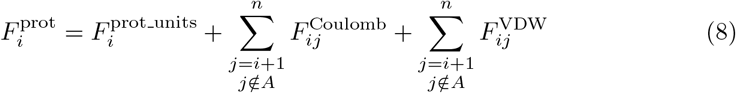

where *i* and *j* denote atoms in the protein, *A* denotes the fragment atom *i* belongs to. AI^2^BMD offers a significant advancement over QM-MM methods by extending ab initio calculations from a limited quantum mechanical region to the protein without prior knowledge.

### ViSNet-PIMA

We adopt a two-scale design. The ViSNet layers (cutoff *r*_*c*_ = 5 Å) learn rich local features, while the Physics-Informed Multipole Aggregator (PIMA) layers learn non-local electrostatic polarization with a large influence radius (e.g., *R* = 15 Å). Concretely, PIMA maintains learnable charge embeddings *q*_*i*_ and dipoles ***µ***_*i*_, and computes charge– charge, charge–dipole, and dipole–dipole interactions with analytic kernels, followed by a small number of differentiable polarization updates. This physics-informed factorization makes PIMA considerably more both parameter-efficient and data-efficient than a brute-force large-cutoff GNN, while preserving end-to-end energy conservation (forces analytic gradients of the total energy).

ViSNet-PIMA takes the atomic numbers and coordinates as inputs. These inputs are first encoded into the invariant scalar features 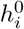 and the equivariant vectorized features 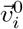 through the embedding block. Then these features are updated in the stacked local interaction blocks, where local atomic environments are encoded via the vector-scalar interactive message passing (ViS-MP) mechanism. For more information about ViSNet layers, please refer to ViSNet[13] for details. The final outputs of the ViSNet layers are the refined scalar features 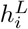 and vector features 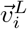, obtained after processing through all *l*_max_ layers (denoted as *L* layers for simplicity). The updated invariant and equivariant features are then fed into PIMA blocks to encode the non-local interactions in the molecule. PIMA starts by transforming the equivariant features 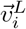 into initial values of dipole embeddings 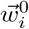 based on the output invariant features 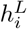 from the ViSNet layers.The dipole embbedings are learnable high-dimensional vectors that serve as internal inputs to the PIMA module.

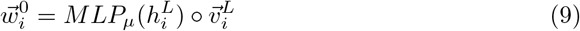

*h*^*L*^ ∈ *R*^*D*^, 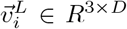, where D demotes the dimension of hidden channels. *MLP*_*µ*_ detnotes the non-linear transformation of 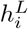, and its Hadamard product with 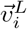 mirrors the definition of dipole moment ∫_*V*_ **r***ρ*(**r**)*d* **r** in multipole expansion.

After obtaining the initial values of dipole embeddings 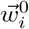, PIMA learns and represents the molecular non-local interactions by two phases. The “Field Response” phase is for dynamic modeling of molecular dipoles’ polarization responses to the external electric fields, where atomic dipole embeddings 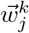 are iteratively updated through environmental modulation till convergence. The following “Interaction Integration” phase integrates these converged dipole interactions with local chemical environments, further enhancing atomic features 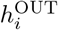 for subsequent energy prediction via message aggregation.

Specifically, there are two implementations of PIMA, which differ in whether to take the information of point charges into account. In the first kind of implementation, only the interactions among dipole embeddings are considered. For the (*L*+*k*)-th layer (corresponding to the *k*-th PIMA layer), the equivariant messages are calculated as follows:

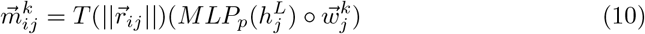

where *MLP*_*p*_ controls the intensity of the dipole embedding of atom *j* when inducing the dipole embedding of atom *i*. 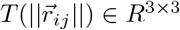 is the interaction matrix of dipole-dipole interactions, which is calculated in the following formula:

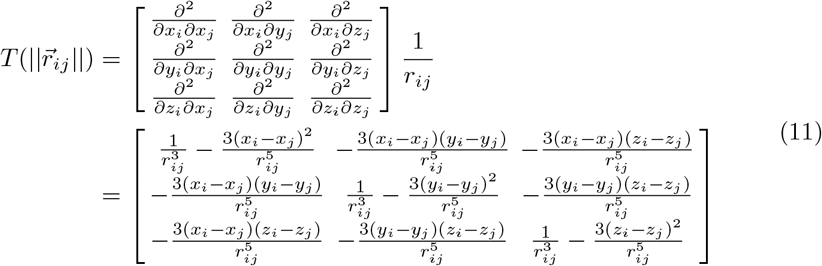

After calculating 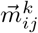, the dipole embedding of atom *i* is updated as follows:

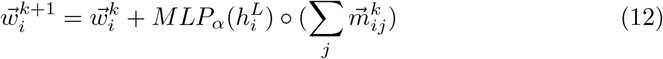

where *MLP*_*α*_ can be interpreted as the transformation of invariant features into polarizability, and *j* belongs to any atoms within a threshold except atom *i*.

In the “Interaction Integration” phase, the invariant features are updated via the following formula:

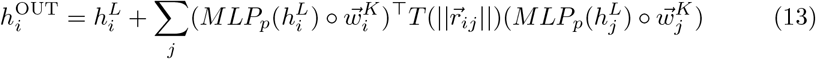

which incorporates non-local dipole-dipole interactions into invariant features.

The other implementation of PIMA additionally models the charge-dipole and charge-charge interactions. Besides dipole embeddings, we further defined the charge embeddings as follows:

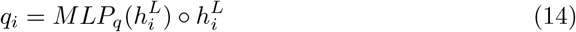

Since the dipole moments instead of the point charges can be polarized in polarizable force fields[52, 53], we keep the charge embeddings fixed during the “Field Response” phase. The charge embbedings are also learnable high-dimensional features that serve as internal inputs to the PIMA module. When taking the charge embeddings into account, the equivariant messages in the *k*-th PIMA layer are calculated as follows:

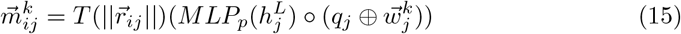

where *q*_*j*_ and 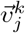 are concatenated together, resulting in a new vector in *R*^4*×D*^. 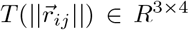 is the interaction matrix of dipole-dipole interactions and dipole-charge interactions, which is calculated in the following formula:

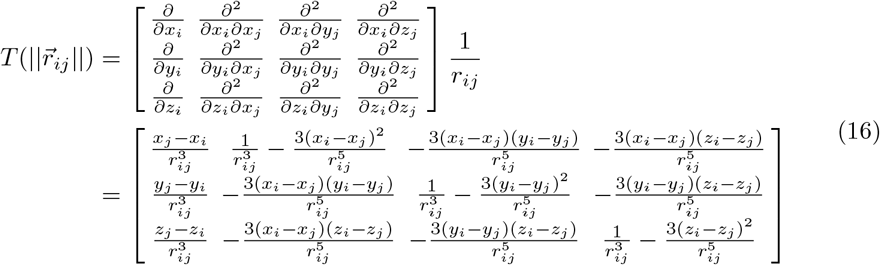

Similarly, the dipole embeddings are updated in Equation 12.

In the “Interaction Integration” Phase, both the charge embedding *q*_*i*_ and the dipole embedding 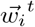 are involved:

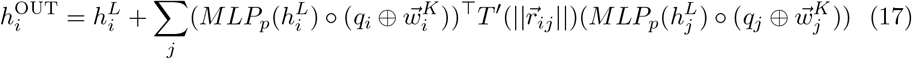

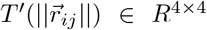 is the interaction matrix of dipole-dipole interactions, dipole-charge interactions and charge-charge interactions, which is calculated in the following formula:

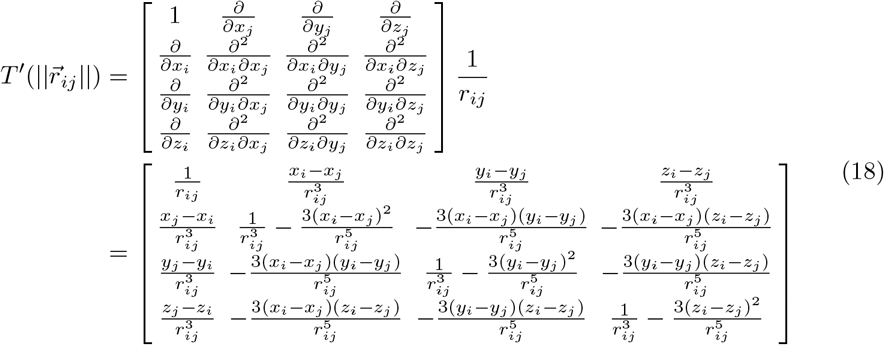

In our evaluations, the two kinds of implementations are both effective. Specifically, the first kind of implementation is more efficient, while the other one is more expressive. It depends on the specific tasks to determine which kind of implementation to be used. For example, in this work, the first implementation was used for model training on MD22 and AIMD-Chig datasets, while the second implementation was used for the dimer dataset and the integration with AI^2^BMD.

After obtaining the updated invariant features and equivariant features, these features are transformed into atomic energy *E*_*i*_ in the output block following the operations in ViSNet[13]. The total energy is calculated as the sum of atomic energies and the atomic forces are calculated as the derivative of energy with respect to atomic coordinates:

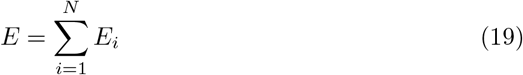

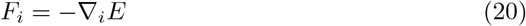

### Incorporation ViSNet-PIMA into AI^2^BMD

The input protein first undergoes the AI^2^BMD fragmentation scheme, which splits the protein into *n* dipeptides and *n* −1 overlapping ACE-NMEs units. These fragments are then fed into the ViSNet trained on PUD dataset. The intra-fragments interactions are then calculated based on each atom’s embedding of its local environment (*h*_*i*_ and 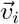) by ViSNet. For each fragment, its energy and atomic forces are calculated, and these intra-fragments interactions are combined into *E*^prot_units^ and 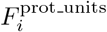. As for the inter-fragments interactions, instead of adopting the classical Coulomb and Lennard-Jones potential, we utilize ViSNet-PIMA to learn these interactions. Instead of training the ViSNet-PIMA model from scratch, we transfer the knowledge learned by the intra-fragment ViSNet into ViSNet-PIMA.Supplementary Table S5 presents a direct comparison between the baseline AI^2^BMD and the proposed AI^2^BMD-PIMA. The embeddings learned within each fragment are first combined as follows:

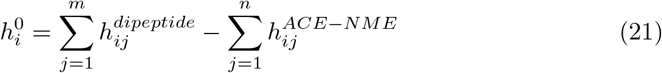

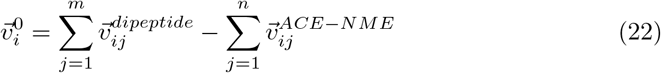

where *m* represents all of the dipeptides the atom *i* belongs to, *n* represents all of the ACE-NME fragments the atom *i* belongs to, and *h*_*ij*_ and 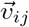 represent atom *i*’s embeddings calculated within the *j*-th protein unit.

Through this combination scheme, each atom of the protein is initially assigned with an invariant feature 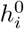 and an equivariant feature 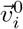, which represent its local environment based on the intra-fragment ViSNet. These features are then fed into ViSNet-PIMA to learn the non-local inter-fragment energy *E*^*inter_frag*^ and force 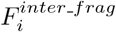. The energy and forces of the whole protein are calculated as a combination of local intra-fragments interactions and non-local inter-fragments interactions:

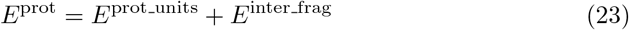

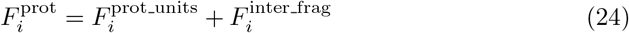

### “Transfer Learning-Pretraining-Finetuning” scheme

In AI^2^BMD, ViSNet is initially trained on protein units to effectively capture local molecular knowledge across various chemical environments. The training process leverages extensive datasets that encompass a wide variety of protein structures, enabling the model to generate continuous latent atom embeddings. These embeddings describe local chemical environments derived from the molecular graph, allowing for a detailed representation of interactions at the atomic level. These atom embeddings are then transformed into equivariant and invariant features that preserve appropriate local chemical symmetries and geometries as demonstrated in the above section.

Subsequently, ViSNet-PIMA harnesses the transferred knowledge to learn the nuances of Coulomb and van der Waals interactions on a pretraining dataset. The pretraining phase is essential, as it allows the model to establish a fundamental understanding of local and non-local interactions informed by a diverse array of protein configurations. The dataset utilized in this phase is derived from simulation trajectories conducted with AI^2^BMD across various protein structures, ensuring that it captures a wide range of conformational states. By sampling from these exhaustive simulations, the dataset provides rich and varied examples of protein behavior, enabling the model to learn the underlying principles governing local and non-local interactions. To achieve precise interaction calculations, we incorporate a limited number of DFT calculations during the finetuning phase. In this stage, the model is refined using a sampled DFT dataset to integrate high-level quantum mechanical principles, thereby enhancing its ability to predict complex non-local and local interactomic interactions.

### Experimental settings

#### Implementation

The experiments of training models on the MD22, Chignolin, dimer dataset and the AI^2^BMD-PIMA are implemented by PyTorch 1.11.0 with 4 NVIDIA GPU cards. To stabilize and accerate MD simulations, the experiments on the solvated system and the separate ion compounds are conducted using a JAX implementation of ViSNet-PIMA. The training and simulation of the JAX-version ViSNet-PIMA are based on the MLIP framework [54]. We construct the loss function to the total energy *E* as well as the forces *F* ^*j*^ for each atom.

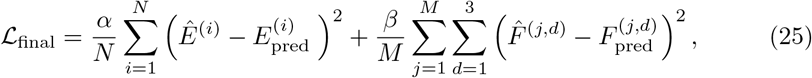

where *N* is the number of samples, *M* is the number of force samples, which is typically related to the number of atoms. *α* and *β* are the weights used to balance the energy loss and force loss.

#### Hyperparameters

For the MD22 and AIMD-Chig datasets, we set the weight *α* of the energy loss to 0.05 and the weight *β* of the force loss to 0.95. The batch size was selected from the values 2, 4, and 8 respectively, based on GPU memory constraints. The learning rate varied from 1e-4 to 4e-4 for different kinds of molecules. The neighboring atom threshold for relatively small molecules was set to 5Å; while for larger molecules, it was set to 4Å. We employed a patience of 5 epochs for early stopping, with a learning rate decay factor of 0.8. For the dimer dataset, we chose ViSNet-PIMA model with the threshold of 3Å, two ViSNet blocks and three PIMA blocks. In the experiments for AI^2^BMD-PIMA, intra-fragment interactions were modeled using a 6-layer ViSNet with a cutoff of 5Å, as described in [12]. Inter-fragment interactions were learned from ViSNet-PIMA, which consists of one ViSNet layer and three PIMA layers. The weight *α* for the energy loss was set to 1, while the weight *β* for the force loss was set to 100. The batch size was configured to 2, and the initial learning rate was set to 1e-4. Additional details regarding the hyperparameters of models are provided in theSupplementary Table S7-S10.

#### Dataset settings

The AIMD-Chig dataset contains 2 million distinct conformations of the 166-atom mini-protein Chignolin, capturing a wide variety of conformational modes. We selected the 5% subset of AIMD-Chig, which contained 100 thousand conformations of Chignolin for our evaluations in this study. The data split of Chignolin was chosen to be consistent with [35]. For the MD22 dataset, we use the same number of molecules as in [13] for training and validation, and the rest as the test set. For the dimer dataset, we adopted the structures and energies calculated at DFT level from SPICE dataset[40, 41]. 80 % pairs of dimers were kept for training and validation, and the rest 20 % were kept for evaluation. For the data generation utilized in training AI^2^BMD-PIMA, we initiated simulations from 20 distinct conformations of four proteins (Protein Data Bank (PDB) IDs: chignolin, 5AWL; Trp-cage, 2JOF; WW domain, 2F21; ABD, 1PRB). These simulations were conducted for 100 ps at 300 K with a timestep of 1 fs, driven by AI^2^BMD. Conformations were sampled every 100 fs from each trajectory to serve as the training data for the pretraining phase, resulting in a dataset comprising over 80,000 data points, each annotated with energy and force values by AI^2^BMD. For the finetuning phase, we randomly selected approximately 50 conformations from each trajectory and computed their energy and forces at the M06-2X/6-31G* level[55] using ORCA 5.0.1 software[56], resulting in approximately 1,000 data points per protein, each labeled with DFT-calculated energy and forces.

#### MD simulation settings

For the MD simulation settings on Ac-Ala3-NHMe, stachyose, buckyball catcher in the MD22 dataset and the separate ion compounds, the simulations were performed at 300 K using Langevin thermostat with a friction coefficient of 0.01 *fs*^−1^ (10 *ps*^−1^) under the ASE framework[45]. We used ViSNet-PIMA to run the simulations with a timestep of 1 fs. To evaluate the relative fluctuations, we chose one conformation per picoseconds along the trajectories and calculated its potential energy at PBE-d3bj/6-31G* level using ORCA 5.0.1. The MD simulations of solvated p-Cresol were conducted using the umbrella sampling method across 15 windows. A harmonic biasing potential with a spring constant of 0.1 eV/^Å2^ was applied to restrain the distance between the atoms closest to the centroid of each p-Cresol monomer. The window centers were spaced along this reaction coordinate, ranging from 10 Å to 2 Å. Each window was simulated for a duration of 10 ps, and the final Potential of Mean Force (PMF) was calculated using the pymbar library [57].

## Data availability

The SPICE, MD22 and AIMD-Chig datasets are publicly available (see “Methods” for more details). Other datasets used in this work (Chignolin/Trp-cage/WW/ABD), source data files underlying all main figures and MD simulation trajectories will be available at https://wanggroup.ai/software upon publication of the manuscript.

## Code availability

The source code for reproducing the findings in this paper will be available at https://wanggroup.ai/software upon publication of the manuscript.

## Author contributions

T. W. led, conceived, designed and supervised this study. T. C. and W. Z. did this study when they did internships at T. W.’s lab. T. C. and W. Z. trained and analyzed all machine learning models on MD22 and AIMD-Chig datasets. Z. W. performed and analyzed the simulations on MD22 molecules. T. C. and Z. W. trained and analyzed the models on dimer dataset. Z. W. performed and analyzed AI2BMD simulations with ViSNet-PIMA. T. W., T. C. and W. Z. wrote the manuscript. All authors approved the final version of the manuscript.

