## Supplementary Tables and Figures for "Enhancing non-local interaction modeling for ab initio biomolecular calculations and simulations with ViSNet-PIMA"

#### List of Tables


S1 Mean absolute errors (MAEs) of energy (kcal/mol) and force (kcal/(mol·Å)) predictions on AIMD-Chig dataset. The best one in each category is highlighted in bold. . . . . . . 1

S2 Ablation study of the PIMA module on double-walled nanotube in MD22 dataset. Mean absolute errors (MAE) of energy (kcal/mol) and force (kcal/(mol·Å)) are evaluated. The "Total" column represents the total number of interaction layers, including both ViSNet layers and PIMA layers. "ViSNet-L layer" denotes ViSNet layer with a cutoff of 100Å for neighboring atoms. The "Order" column specifies whether the sequence is in normal order (ViSNet+PIMA) or reverse order (PIMA+ViSNet). The "PIMA-S" column represents a PIMA variant that updates both equivariant and invariant features simultaneously during the "Field Response" phase. "OOM" denotes the model runs out of memory on a A100 GPU with 80GB memory. . . . . . . 2

S3 Training time (second per epoch (s/epoch)) for Ac-$Ala_3$-NHMe on a A800 GPU. . . . . . . 3

S4 Comparison of model size, training cost and prediction accuracy on AT-AT-CG-CG in MD22 dataset. The model training time was estimated on a A800 GPU. . . . . . . 4

S5 Comparison between $AI^2BMD$ and $AI^2BMD$-PIMA. . . . . . . 5

S6 The MD simulation speed (second/100 steps) for the protein Trpcage on a A800 GPU. . . . . . . 6

S7 Hyperparameters for ViSNet-PIMA trained on MD22 dataset. . . . . . . 7

S8 Hyperparameters for ViSNet-PIMA trained on AIMD-Chig dataset. . . 8

S9 Hyperparameters for ViSNet-PIMA trained on dimer dataset. . . . . 9

S10 Hyperparameters for $AI^2BMD$-PIMA pretraining and finetuning processes on proteins. . . . . . . 10

### List of Figures


S1 **Evaluations of non-local interactions on the dimer dataset by ViSNet-PIMA. a-f.** Examples of binding energy curves of different dimers predicted by DFT, ViSNet-PIMA and ViSNet, respectively. . 11

S2 **Relative errors of energy calculated by ViSNet and ViSNet-PIMA on dimer systems. a.** a polar compound and a cation. **b.** a polar compound and an anion. **c.** a polar compound and a charged compound. **d.** two polar compounds. The systems in a to d correspond to those in Figure 3 of the manuscript. . . . . . . . . . . . . . . . . . . . 12

S3 **Comparisons of ViSNet-PIMA and ViSNet with MM corrections of empirical Coulomb and Van der Waals functions (termed as "ViSNet-MM"). a.** The binding energy curves of a dimer between a polar compound and a cation (sodium ion) calculated by DFT, ViSNet, ViSNet-PIMA and ViSNet-MM. **b.** The binding energy curves of a dimer between a polar compound and an anion (chloride ion) calculated by the four methods. **c.** The binding energy curves of a dimer between a polar compound and a charged compound calculated by the four methods. **d.** The binding energy curves of a dimer between two polar compounds calculated by the four methods. . . . . . . . . . 13

S4 **Validation of energy and force consistency in MD simulations under NVE ensemble driven by ViSNet-PIMA. a.** Total energy of Ac-Ala$_3$-NHMe remains conserved during a 10 ps NVE simulation driven by the ViSNet-PIMA. **b–d.** Cartesian force components fluctuate around zero, confirming force consistency with the conserved energy. . 14

S5 **Evaluations of MD simulations for Ac-Ala$_3$-NHMe, stachyose, buckyball catcher driven by ViSNet-PIMA. a-c.**The visualization of initial structures of Ac-Ala3-NHMe, stachyose and buckyball catcher, respectively. **d-f.** Relative energy fluctuations of the 100 ps simulation trajectories calculated by ViSNet-PIMA and DFT. Results of ViSNet-PIMA are plotted with a solid blue line, while those of DFT are plotted with a dashed orange line. The relative energy has that of the initial structure subtracted. **g-i.** Interatomic distance distributions predicted by ViSNet-PIMA. **j-l.** Vibrational spectrums calculated as the Fourier transform of the velocity auto-correlation function. . . . . . . . . . . . 15

S6 **Simualations of two separate charged compounds by MACE-OFF and ViSNet-PIMA. a-b.** Representative organic cation–anion ion pairs, shown at the initial separation of 12 Å (**a**) and after approaching into close contact (**b**). **c–e.** Simulation trajectories sampled by MACE-OFF, where the limited receptive field leads to progressive dissociation of the two ions. **f–h.** Simulation trajectories sampled by ViSNet-PIMA, in which long-range interactions are properly captured, driving the ions to approach each other and stabilize at an equilibrium distance. . . . . . . . . . . . . . . . . . . . . . . . . . . . . . . . . . . 16

S7 **Performance evaluation on water cluster test subset in SPICE dataset between MACE OFF(L) and ViSNet-PIMA.** . . . . . . 17

S8 **The potential of mean force (PMF) profile of the association of two solvated p-Cresol molecules from the simulations driven by ViSNet-PIMA with explicit solvent.** The minimum corresponds to the stable T-shape structure of the dimer system. . . . . . . . . . . 18
S9 **Distributions of energy and force on the pretraining dataset of AI$^2$BMD with ViSNet-PIMA.** The energy distributions of Chignolin, Trp-cage, WW and ABD are shown in **a**, **c**, **e** and **g** respectively, while the force distributions of these proteins are shown in **b**, **d**, **f** and **h** respectively. . . . . . . . . . . . . . . . . . . . . . . . . . . . . . . . . . . 19
S10 **Distributions of energy and force on the finetuning dataset of AI$^2$BMD with ViSNet-PIMA.** The energy distributions of Chignolin, Trp-cage, WW and ABD are shown in **a**, **c**, **e** and **g** respectively, while the force distributions of these proteins are shown in **b**, **d**, **f** and **h** respectively. . . . . . . . . . . . . . . . . . . . . . . . . . . . . . . . . . . 20
S11 **Evaluations on energy calculations for folded, intermediate and unfolded structures of different proteins. a.** The MAE of energies predicted by the vanilla AI$^2$BMD. **b.** The MAE of energies predicted by AI$^2$BMD with ViSNet-PIMA. . . . . . . . . . . . . . . . . . . . . . . 21
S12 **Distributions of the residual atomic forces for folded, intermediate and unfolded structures of Chignolin (a), Trp-cage (b), WW (c) and ABD (d), respectively.** The orange, green and purple shadows represent the results of folded, intermediate and unfolded structures, respectively. . . . . . . . . . . . . . . . . . . . . . . . . . . 22
S13 **Comparison of time consumption of energy calculation for four proteins.** DFT calculations were carried out on a GPU. . . . . . . . . 23

**Table S1** Mean absolute errors (MAEs) of energy (kcal/mol) and force (kcal/(mol·Å)) predictions on AIMD-Chig dataset. The best one in each category is highlighted in bold.

| | PaiNN | SO3krates | Equiformer v2 | Mace | ViSNet | ViSNet-PIMA |
|---|---|---|---|---|---|---|
| Energy | 3.6935 | 5.5264 | 4.8043 | 3.8241 | 3.8174 | **1.9841** |
| Force | 0.6701 | 0.5995 | 0.5338 | 0.5553 | 0.5516 | **0.4798** |

**Table S2** Ablation study of the PIMA module on double-walled nanotube in MD22 dataset. Mean absolute errors (MAE) of energy (kcal/mol) and force (kcal/(mol·Å)) are evaluated. The "Total" column represents the total number of interaction layers, including both ViSNet layers and PIMA layers. "ViSNet-L layer" denotes ViSNet layer with a cutoff of 100Å for neighboring atoms. The "Order" column specifies whether the sequence is in normal order (ViSNet+PIMA) or reverse order (PIMA+ViSNet). The "PIMA-S" column represents a PIMA variant that updates both equivariant and invariant features simultaneously during the "Field Response" phase. "OOM" denotes the model runs out of memory on a A100 GPU with 80GB memory.

| #Total layer | #ViSNet layer | #PIMA layer | #ViSNet-L layer | Order | PIMA-S | Energy | Force |
|---|---|---|---|---|---|---|---|
| 6 | 6 | 0 | 0 | ✓ | ✗ | 0.8002 | 0.3621 |
| 6 | 9 | 0 | 0 | ✓ | ✗ | 0.7449 | 0.2489 |
| 9 | 6 | 0 | 3 | ✓ | ✗ | 0.7665 | 0.2832 |
| 9 | 0 | 0 | 6 | ✓ | ✗ | OOM | OOM |
| 9 | 6 | 3 | 0 | ✗ | ✗ | 0.8239 | 0.3144 |
| 9 | 6 | 3 | 0 | ✓ | ✓ | 0.7734 | 0.2444 |
| 9 | 6 | 3 | 0 | ✓ | ✗ | **0.4522** | **0.2243** |

**Table S3** Training time (second per epoch (s/epoch)) for Ac-$Ala_3$-NHMe on a A800 GPU.

| Model | ViSNet | ViSNet (large) | ViSNet-PIMA | MACE |
|---|---|---|---|---|
| Total layer | 6 | 9 | 9 | 2 |
| ViSNet layer | 6 | 9 | 6 | - |
| PIMA layer | - | - | 3 | - |
| MACE layer | - | - | - | 2 |
| Time (s/epoch) | 110 | 159 | 132 | 1,024 |

**Table S4** Comparison of model size, training cost and prediction accuracy on AT-AT-CG-CG in MD22 dataset. The model training time was estimated on a A800 GPU.

| **Item** | ViSNet (small) | ViSNet-PIMA (small) | PaiNN | MACE |
|---|---|---|---|---|
| Number of parameters (million) | 3.3 | 3.8 | 3.2 | 1.0 |
| Training time per epoch (s) | 25 | 28 | 16 | 302 |
| Force MAE (kcal/(mol*Å)) | 0.16 | 0.13 | 0.35 | 0.15 |

**Table S5** Comparison between AI$^2$BMD and AI$^2$BMD-PIMA.

| Item | AI$^2$BMD | AI$^2$BMD-PIMA |
|---|---|---|
| Intra-fragment (local) modeling | ViSNet (EGNN on fragments) | ViSNet (EGNN on fragments) |
| Inter-fragment (non-local) modeling | Classical MM (Coulomb + vdW) | ViSNet-PIMA (physics-informed MLFF) |
| Intra-fragment learning scheme | Fixed empirical function | Learnt via ViSNet-PIMA with pretraining and finetuning |
| Solvent | Amoeba polarizable force field | Amoeba polarizable force field |
| DFT-level accuracy scope | Intra-fragment-level | Entire molecule (including inter-fragment) |
| Accuracy | Limited by MM approximation | Improved via multipole-informed learning |
| Data efficiency | DFT data required for fragments | DFT data required for fragments and a small finetuning set |

**Table S6** The MD simulation speed (second/100 steps) for the protein Trpcage on a A800 GPU.

| Simulation program | $AI^2BMD$ | $AI^2BMD$-PIMA | MACE-OFF |
|---|---|---|---|
| Time (s/100 steps) | 8 | 10 | 27 |

### Hyperparameters

We show the detailed hyperparameters for different datasets in Table S7, Table S8, Table S9 and Table S10.

**Table S7** Hyperparameters for ViSNet-PIMA trained on MD22 dataset.

| | Ac-Ala3-NHMe | DHA | Stachyose | AT-AT | AT-AT-CG-CG | Buckyball catcher | Double-walled nanotube |
|---|---|---|---|---|---|---|---|
| maximum epochs | 10000 | 10000 | 10000 | 3000 | 3000 | 1000 | 3000 |
| early stopping patience | 600 | 600 | 600 | 600 | 600 | 600 | 600 |
| init learning rate | $4e^{-4}$ | $2e^{-5}$ | $4e^{-4}$ | $2e^{-4}$ | $2e^{-4}$ | $2e^{-4}$ | $8e^{-5}$ |
| lr patience | 30 | 30 | 30 | 30 | 30 | 30 | 30 |
| lr decay factors | 0.8 | 0.8 | 0.8 | 0.8 | 0.8 | 0.8 | 0.8 |
| lr warmup steps | 1000 | 1000 | 1000 | 1000 | 1000 | 1000 | 1000 |
| batch size | 4 | 4 | 4 | 4 | 4 | 4 | 1 |
| hidden size | 256 | 256 | 256 | 256 | 256 | 256 | 256 |
| cutoff | 5.0 | 5.0 | 5.0 | 5.0 | 5.0 | 5.0 | 4.0 |
| forces / energy weight | 0.95/0.05 | 0.95/0.05 | 0.95/0.05 | 0.95/0.05 | 0.95/0.05 | 0.95/0.05 | 0.95/0.05 |

**Table S8** Hyperparameters for ViSNet-PIMA trained on AIMD-Chig dataset.

| Hyperparameters | Search space |
|---|---|
| Number of ViSNet layers | 6 |
| Number of PIMA layers | 3 |
| Hidden size | 256 |
| Cutoff distance | 5.0 |
| Batch size | 4 |
| Initial learning rate | 2e-4 |
| Learning rate patience | 30 |
| Learning rate factors | 0.8 |
| Learning rate warmup steps | 1,000 |
| Learning rate warmup factor | 0.7 |
| Max # of Epochs | 3,000 |

**Table S9** Hyperparameters for ViSNet-PIMA trained on dimer dataset.

| Hyperparameters | Search space |
|---|---|
| Number of ViSNet layers | 2 |
| Number of PIMA layers | 3 |
| Hidden size | 128 |
| Cutoff distance | 3.0 |
| Batch size | 128 |
| Initial learning rate | 1e-4 |
| Learning rate patience | 10 |
| Learning rate factors | 0.8 |
| Learning rate warmup steps | 1,000 |
| Learning rate warmup factor | 0.7 |
| Max # of Epochs | 10,000 |

**Table S10** Hyperparameters for AI$^2$BMD-PIMA pretraining and finetuning processes on proteins.

| Hyperparameters | Search space |
|---|---|
| Number of ViSNet layers | 1 |
| Number of PIMA layers | 3 |
| Hidden size | 128 |
| Cutoff distance | 5.0 |
| Batch size | 2 |
| Initial learning rate | 1e-4 |
| Learning rate patience | 10 |
| Learning rate factors | 0.8 |
| Learning rate warmup steps | 1,000 |
| Learning rate warmup factor | 0.7 |
| Max # of Epochs | 1,000 |

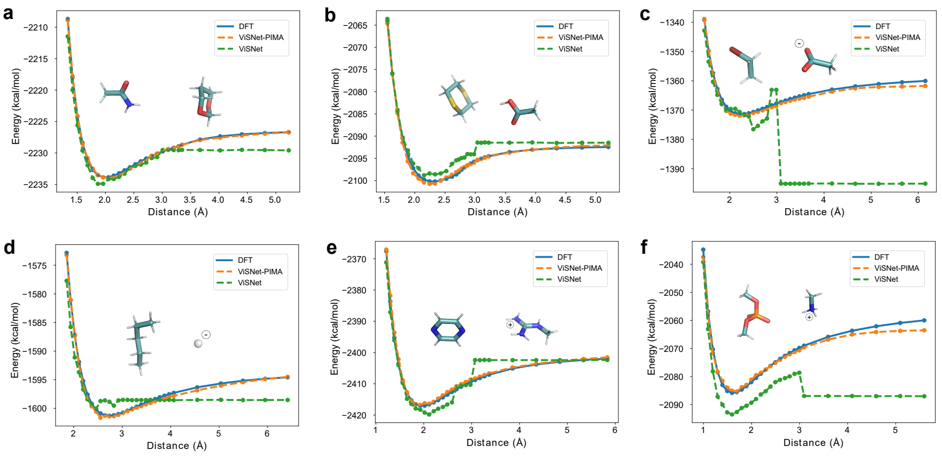


**Figure S1 Evaluations of non-local interactions on the dimer dataset by ViSNet-PIMA. a-f.** Examples of binding energy curves of different dimers predicted by DFT, ViSNet-PIMA and ViSNet, respectively.

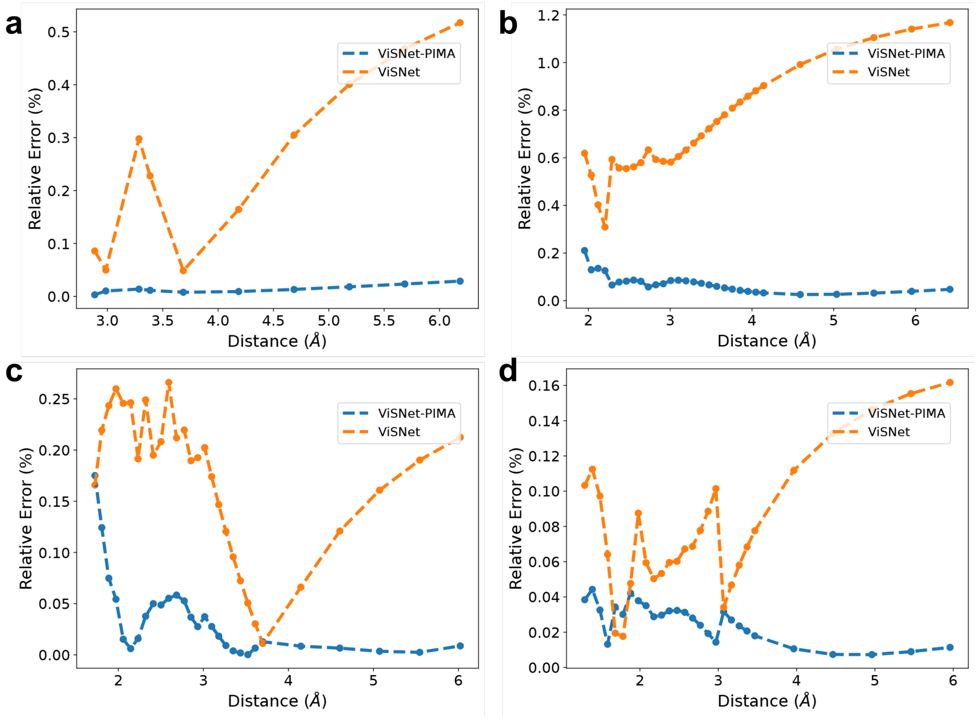


**Figure S2 Relative errors of energy calculated by ViSNet and ViSNet-PIMA on dimer systems.** **a.** a polar compound and a cation. **b.** a polar compound and an anion. **c.** a polar compound and a charged compound. **d.** two polar compounds. The systems in a to d correspond to those in Figure 3 of the manuscript.

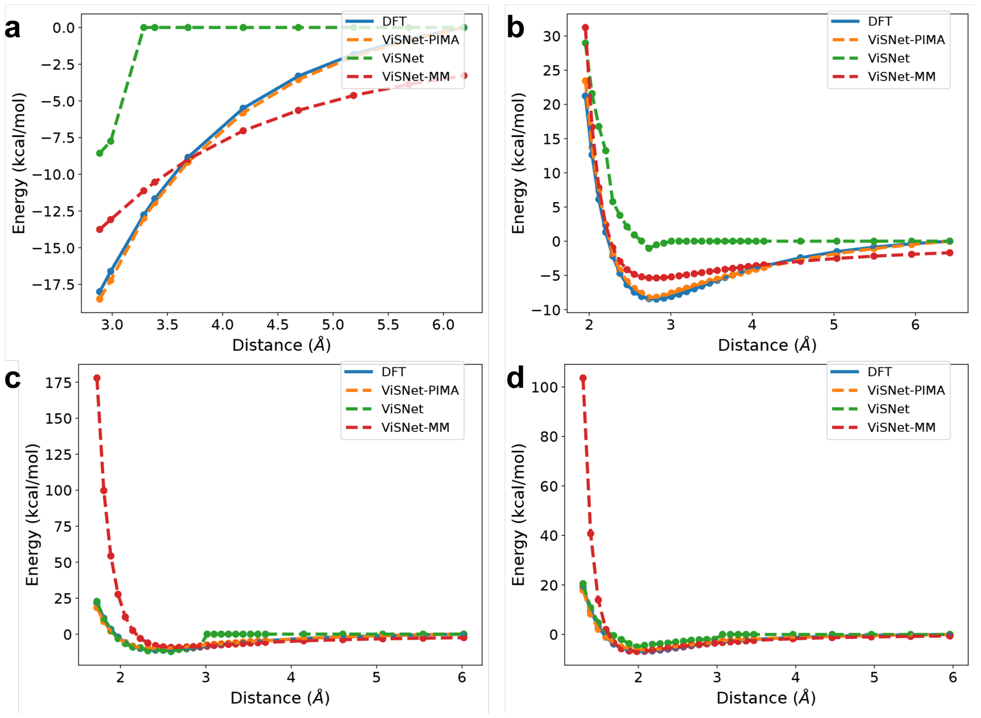


**Figure S3 Comparisons of ViSNet-PIMA and ViSNet with MM corrections of empirical Coulomb and Van der Waals functions (termed as "ViSNet-MM"). a.** The binding energy curves of a dimer between a polar compound and a cation (sodium ion) calculated by DFT, ViSNet, ViSNet-PIMA and ViSNet-MM. **b.** The binding energy curves of a dimer between a polar compound and an anion (chloride ion) calculated by the four methods. **c.** The binding energy curves of a dimer between a polar compound and a charged compound calculated by the four methods. **d.** The binding energy curves of a dimer between two polar compounds calculated by the four methods.

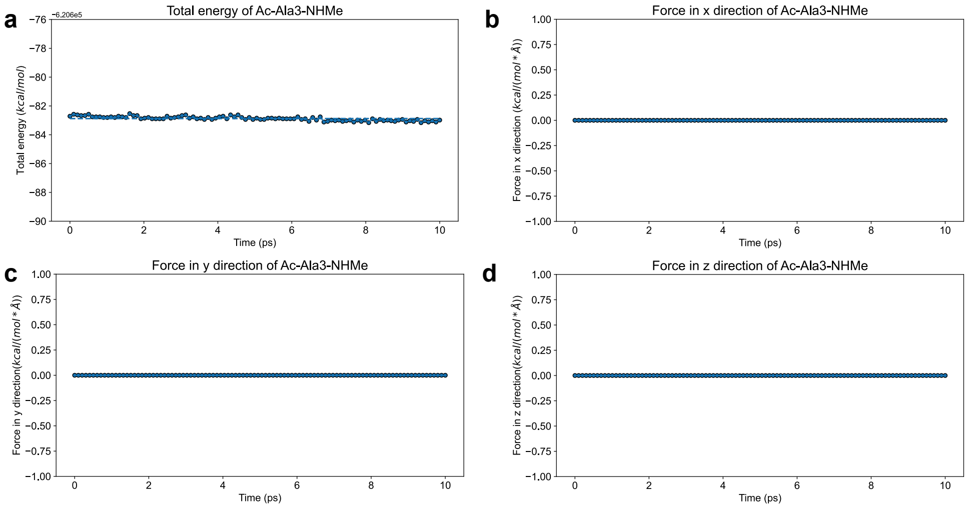


**Figure S4 Validation of energy and force consistency in MD simulations under NVE ensemble driven by ViSNet-PIMA. a.** Total energy of Ac-Ala$_3$-NHMe remains conserved during a 10 ps NVE simulation driven by the ViSNet-PIMA. **b–d.** Cartesian force components fluctuate around zero, confirming force consistency with the conserved energy.

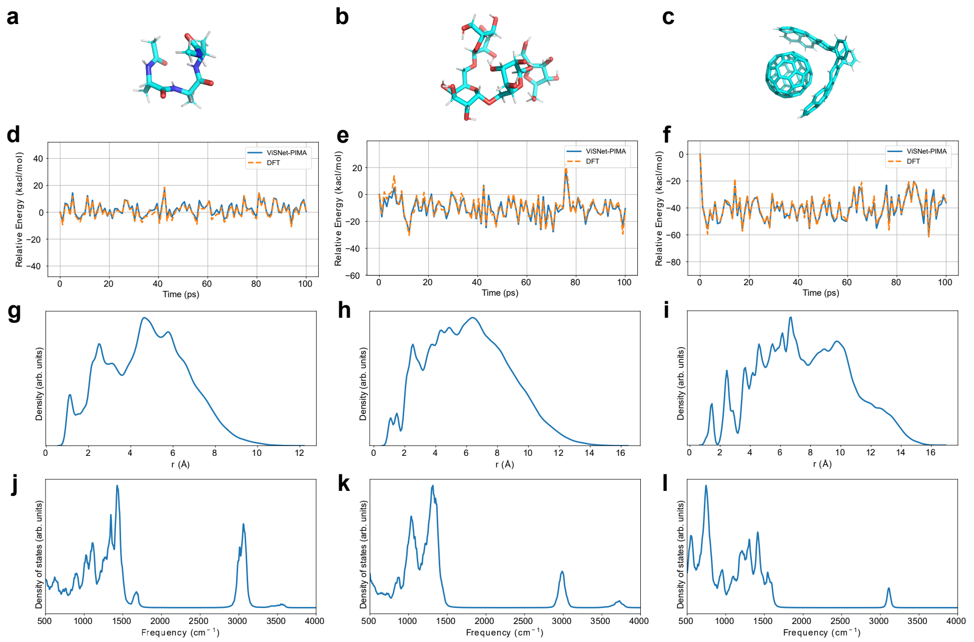


**Figure S5 Evaluations of MD simulations for Ac-Ala$_3$-NHMe, stachyose, buckyball catcher driven by ViSNet-PIMA. a-c.**The visualization of initial structures of Ac-Ala3-NHMe, stachyose and buckyball catcher, respectively. **d-f.** Relative energy fluctuations of the 100 ps simulation trajectories calculated by ViSNet-PIMA and DFT. Results of ViSNet-PIMA are plotted with a solid blue line, while those of DFT are plotted with a dashed orange line. The relative energy has that of the initial structure subtracted. **g-i.** Interatomic distance distributions predicted by ViSNet-PIMA. **j-l.** Vibrational spectrums calculated as the Fourier transform of the velocity auto-correlation function.

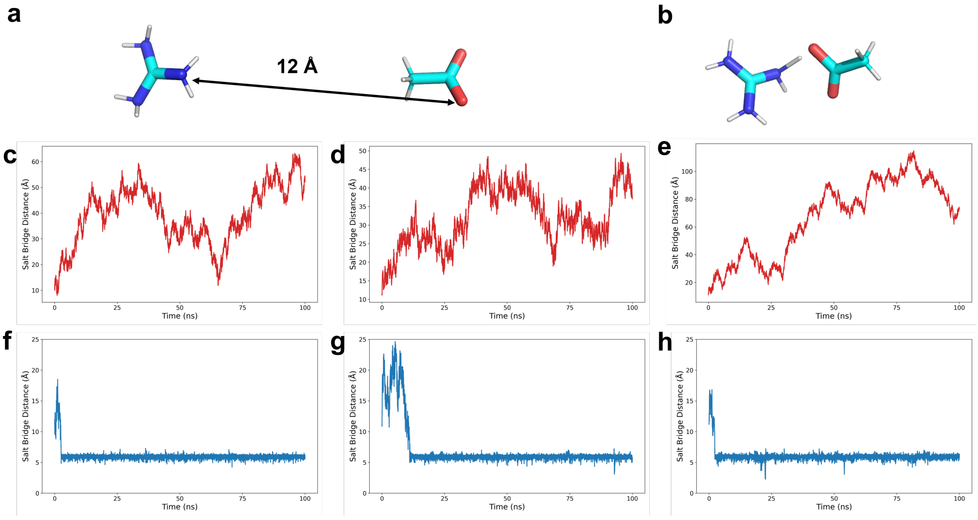


**Figure S6 Simualations of two separate charged compounds by MACE-OFF and ViSNet-PIMA. a-b.** Representative organic cation–anion ion pairs, shown at the initial separation of 12 Å (**a**) and after approaching into close contact (**b**). **c–e.** Simulation trajectories sampled by MACE-OFF, where the limited receptive field leads to progressive dissociation of the two ions. **f–h.** Simulation trajectories sampled by ViSNet-PIMA, in which long-range interactions are properly captured, driving the ions to approach each other and stabilize at an equilibrium distance.

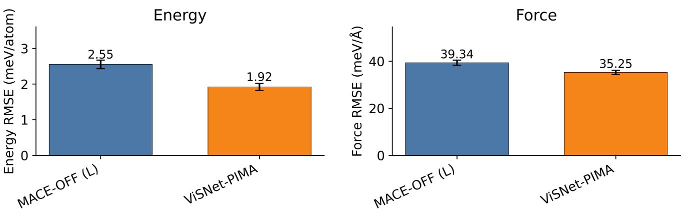


**Figure S7 Performance evaluation on water cluster test subset in SPICE dataset between MACE OFF(L) and ViSNet-PIMA.**

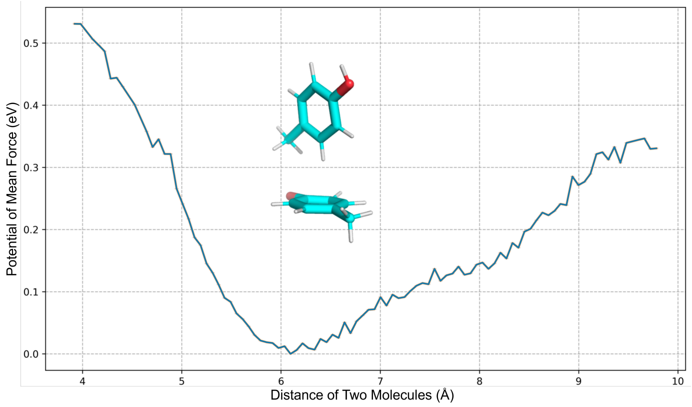


**Figure S8 The potential of mean force (PMF) profile of the association of two solvated p-Cresol molecules from the simulations driven by ViSNet-PIMA with explicit solvent.** The minimum corresponds to the stable T-shape structure of the dimer system.

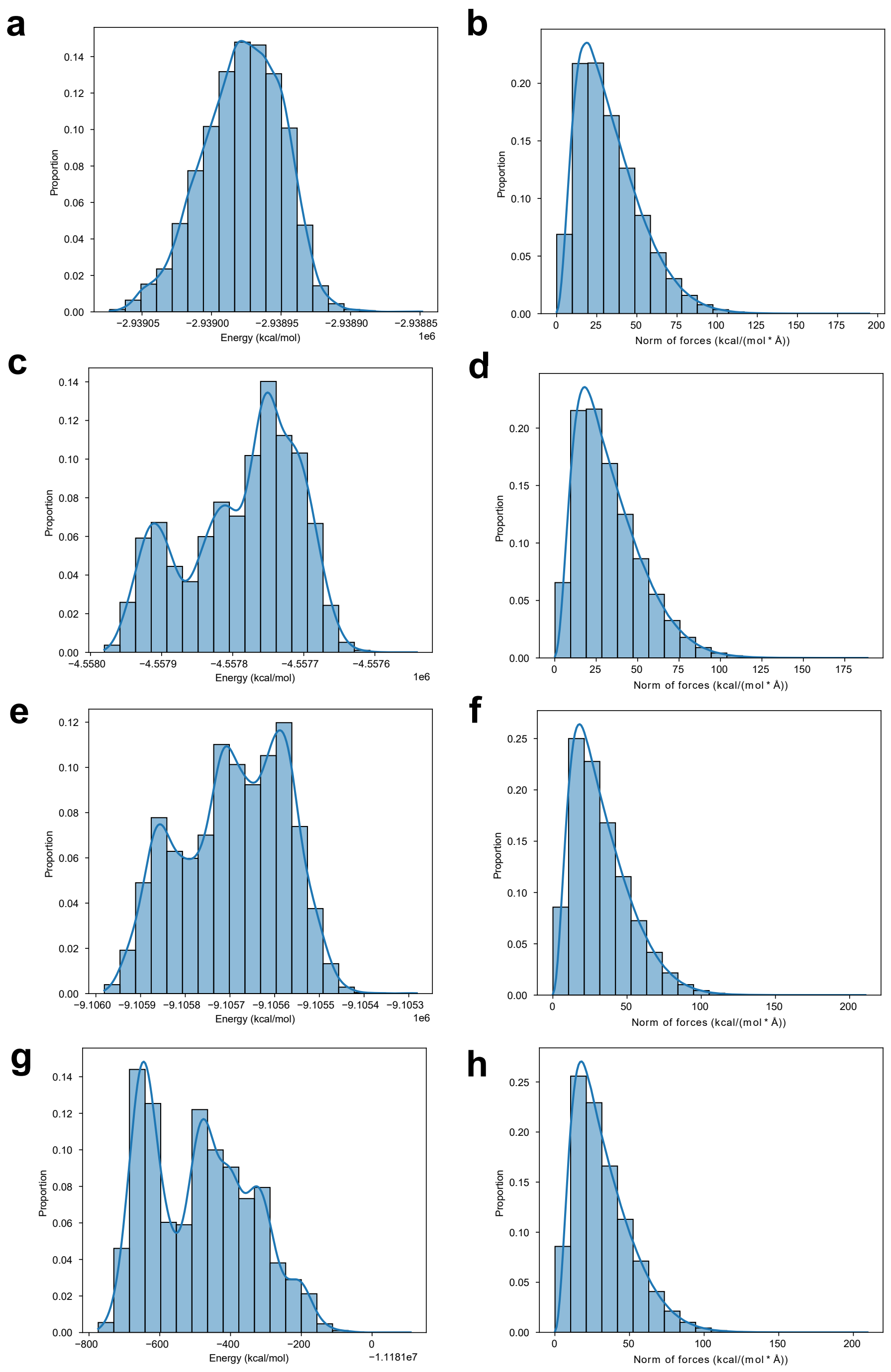


**Figure S9 Distributions of energy and force on the pretraining dataset of AI$^2$BMD with ViSNet-PIMA.** The energy distributions of Chignolin, Trp-cage, WW and ABD are shown in **a**, **c**, **e** and **g** respectively, while the force distributions of these proteins are shown in **b**, **d**, **f** and **h** respectively.

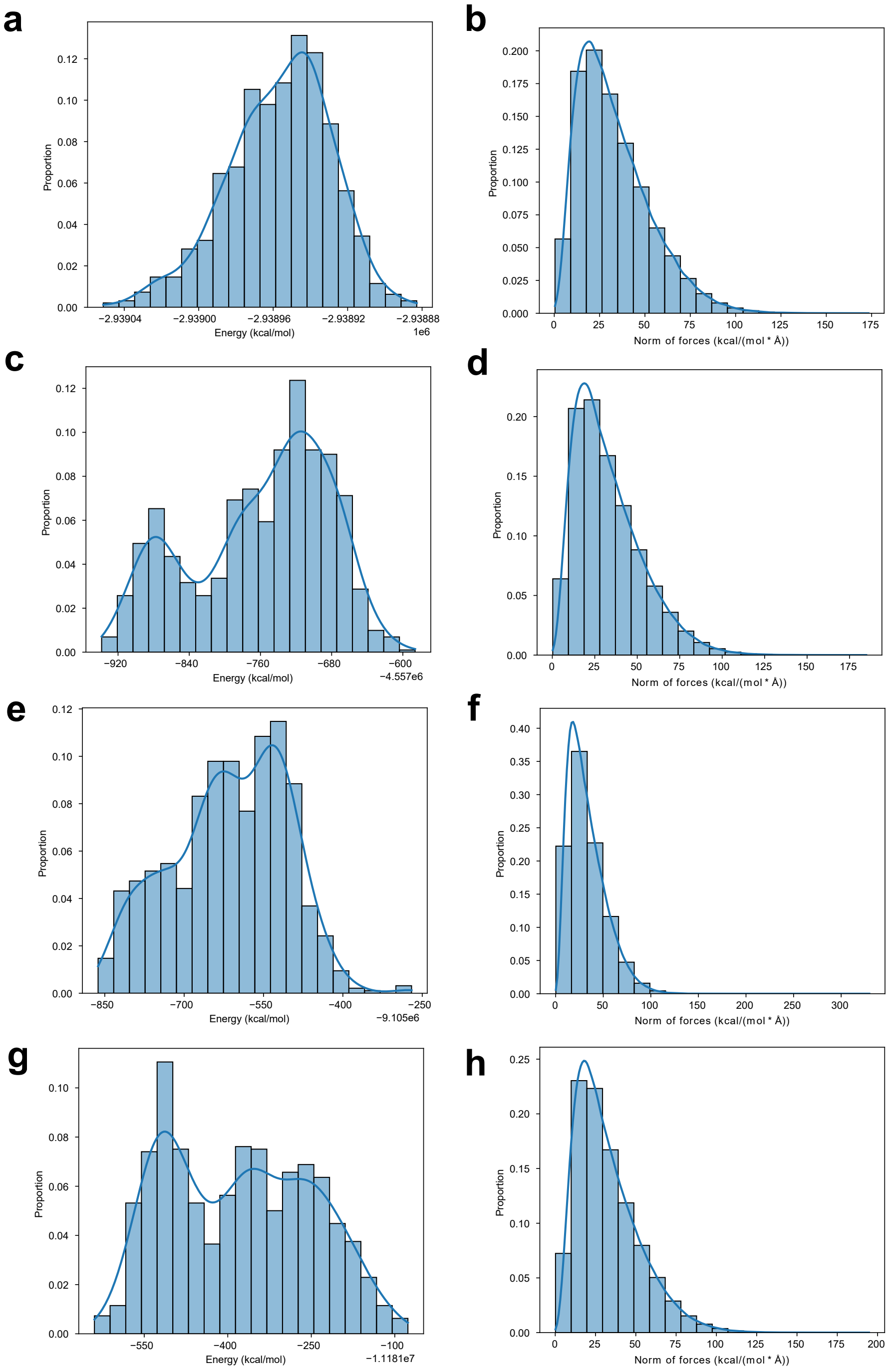


**Figure S10 Distributions of energy and force on the finetuning dataset of AI$^2$BMD with ViSNet-PIMA.** The energy distributions of Chignolin, Trp-cage, WW and ABD are shown in **a**, **c**, **e** and **g** respectively, while the force distributions of these proteins are shown in **b**, **d**, **f** and **h** respectively.

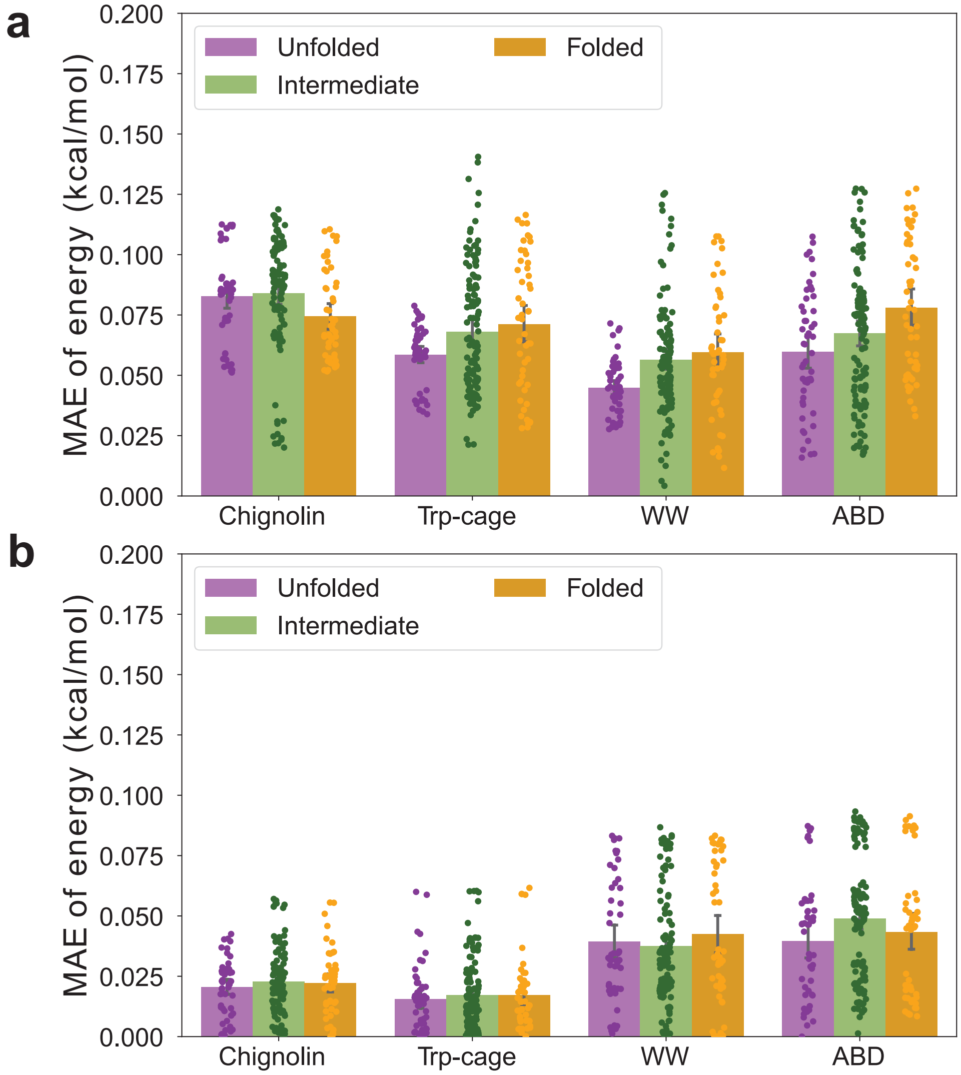


**Figure S11 Evaluations on energy calculations for folded, intermediate and unfolded structures of different proteins. a.** The MAE of energies predicted by the vanilla AI$^2$BMD. **b.** The MAE of energies predicted by AI$^2$BMD with ViSNet-PIMA.

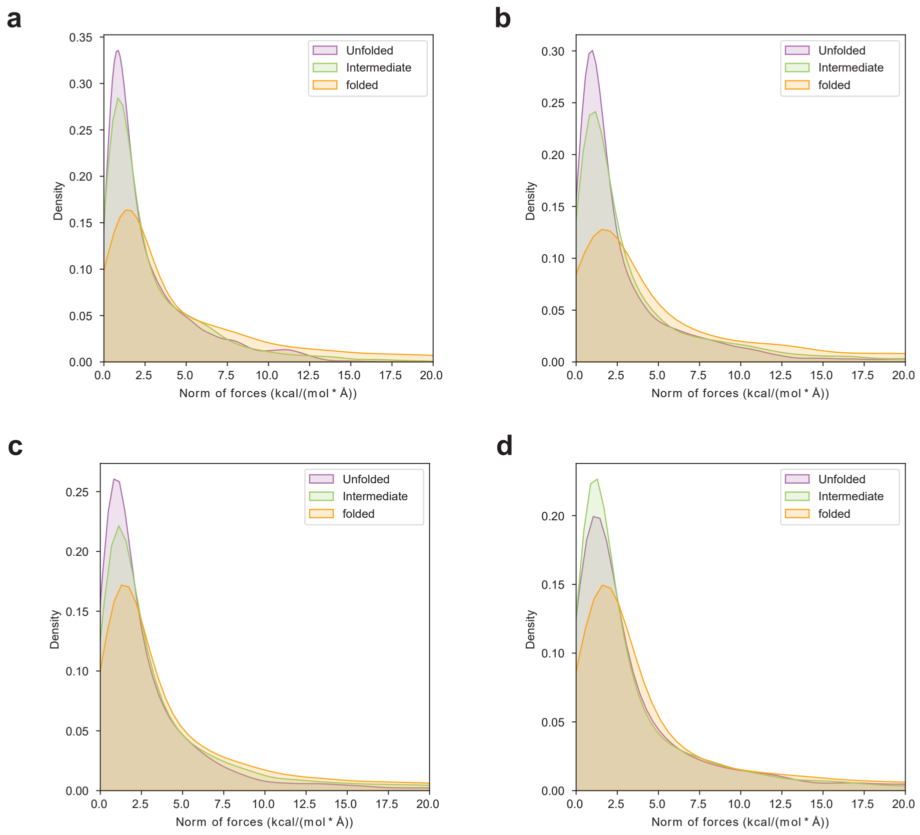


**Figure S12 Distributions of the residual atomic forces for folded, intermediate and unfolded structures of Chignolin (a), Trp-cage (b), WW (c) and ABD (d), respectively.** The orange, green and purple shadows represent the results of folded, intermediate and unfolded structures, respectively.

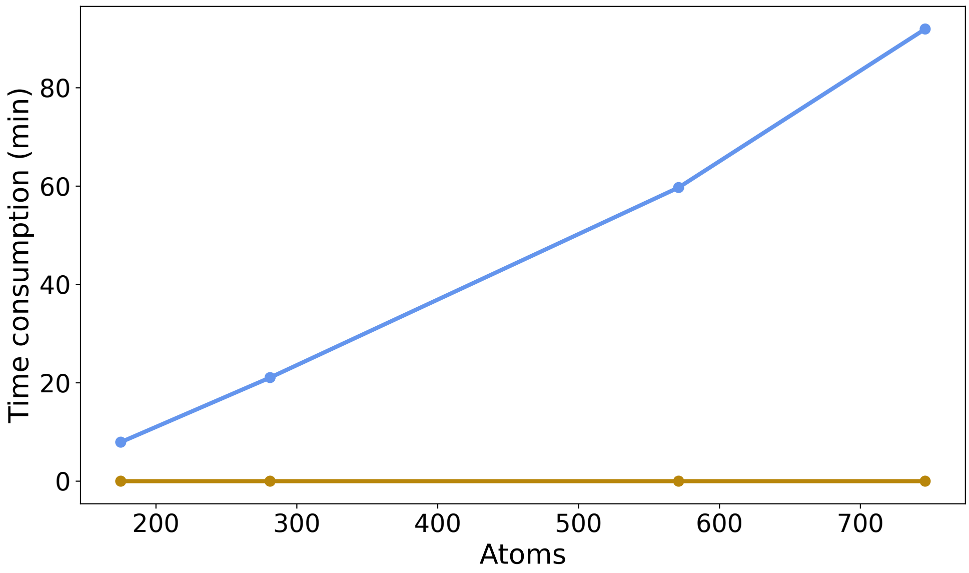


**Figure S13 Comparison of time consumption of energy calculation for four proteins.** DFT calculations were carried out on a GPU.
